# Single-cell transcriptome analysis on the anatomic positional heterogeneity of pig skin

**DOI:** 10.1101/2023.02.17.528908

**Authors:** Qin Zou, Rong Yuan, Yu Zhang, Yifei Wang, Ting Zheng, Rui Shi, Mei Zhang, Yujing Li, Kaixin Fei, Ran Feng, Binyun Pan, Xinyue Zhang, Zhengyin Gong, Li Zhu, Guoqing Tang, Mingzhou Li, Xuewei Li, Yanzhi Jiang

## Abstract

Different anatomic locations of the body skin dermis come from different origins, and its positional hereditary information can be maintained in adults, while highly resolvable cellular specialization is less well characterized in different anatomical regions. Pig is regarded as excellent model for human research in view of its similar physiology to human. In this study, we performed single-cell RNA sequencing of six different anatomical skin regions from the Chenghua pig with superior skin thickness trait. We obtained 215,274 cells, representing seven cell types, among which we primarily characterized the heterogeneity of smooth muscle cells, endothelial cells and fibroblasts. We identified several phenotypes of smooth muscle cell and endothelial cell and presented genes expression of pathways such as the immune response in different skin regions. By comparing differentially expressed fibroblast genes among different skin regions, we considered TNN, COL11A1, and INHBA as candidate genes for facilitating ECM accumulation. These findings of heterogeneity in the main three cell types from different anatomic skin sites will contribute to a better understanding of hereditary information and places the potential focus on skin generation, transmission and transplantation, paving the foundation for human skin priming.

## Introduction

The problem of how hereditary information contributes to anatomical site-specific differences has inspired extensive exploration. The pattern formation of spatial arrangement addresses the expression control of specific genes with a cell type. Anatomical site-specific pattern information is determined in the embryo, and site-specific patterns of cellular specialization could also be maintained throughout adulthood along with continual self-renewal tissues (Rinn et al., 2006). Information on site-specific patterns in anatomical tissue has been uncovered, such as in the heart (Litviňuková et al., 2020) and muscle (De Micheli et al., 2020), but the highly resolvable patterns in cellular specialization are less well understood in physiologically different anatomical skin regions.

Skin is the largest organ, providing physical, chemical and biological barrier for the body. It consists of the upper epidermis and the lower dermis layers separated by the basement membrane, with unambiguous spatial patterns of morphologic and functional specialization (Simpson et al., 2011). Embryological studies have shown that anatomic positional-specific information is provided by the stroma, which is composed of extracellular matrix and mesenchymal or dermal cells during embryogenesis (Rinn et al., 2006). Pioneering studies showed that the different anatomic locations of the body skin dermis arose from different origins. The dorsum dermis originates from the dermato-myotome, the ventral dermis from the lateral plate mesoderm and the face dermis from the neural crest (Jinno et al., 2010; Ohtola et al., 2008; Wong et al., 2006). In adults, dermal cells confer positional identity and memory for skin patterning and function (Driskell and Watt, 2015), raising the question of what regional discrepancy could be maintained against plentiful cellular turnover in skin.

The dermis is composed of resident dermal fibroblasts (FBs), smooth muscle cells (SMCs), endothelial cells (ECs) and immune cells, which provide structure, strength, flexibility, and defense to the skin (Driskell and Watt, 2015). FBs, the main cell type in the dermis, are responsible for the collagen deposits and elastic fibres of the extracellular matrix (ECM) (Parsonage et al., 2005), which are an integral part of skin morphogenesis, homeostasis, and various physiological and pathological mechanisms, including skin development, ageing, healing, and fibrosis (Auxenfans et al., 2009; Driskell et al., 2013; Driskell and Watt, 2015). SMCs, which form blood vessels and arrector pili muscle (APM), play a critical role in controlling blood distribution as well as maintaining the structural integrity of the blood vessels and arrector pili muscle (APM) in skin (Driskell et al., 2013; Liu and Gomez, 2019). ECs organize the vascular plexus, which plays a predominant role in vascular remodeling, metabolism and the immune response in the dermis, and EC metabolism is tightly connected to barrier integrity, immune and cellular crosstalk with smooth muscle cells (Cantelmo et al., 2016; Miyagawa et al., 2019; Tombor et al., 2021). In general, the skin dermal rection is realized by cell-cell communication and dynamic, cell-matrix interactions and regulatory factors.

Given that the pig model is a powerful tool for skin research according to the similar histological, ultrastructural, and physiological functions of skin between humans and pigs, we chose the Chenghua (CH) pig, with its superior skin thickness traits, for investigating regional variation. Here, single-cell RNA sequencing with an unprecedented resolution allows simultaneous profiling of transcriptomes for thousands of individual cells, to focus on six different anatomic sites of CH pig skin tissue. We obtained a single-cell transcriptome atlas of 215,274 cells and identified seven cell types with unique gene expression signatures. In our datasets, we analyzed the three cell types with largest number, including SMCs, ECs and FBs. SMCs revealed the signature of contractile SMCs, mesenchymal-like phenotype and macrophage-like phenotype in healthy skin samples and presented some genes related to ECM-integrins and immune response in different skin anatomic sites. ECs were classified into four EC phenotypes, and the gene expression of integrins, immunity and metabolism across six different anatomic sites of skin was explored. Moreover, comparative differentially expression genes (DEGs) of FBs among different regions showed that TNN, COL11A1, and INHBA might be candidate genes for ECM accumulation. Taken together, the data in this study offer a comprehensive understanding of the single-cell atlas that displays the different skin anatomic sites of pigs, supporting future exploration as a baseline for heathy and morbid human skin.

## Results

### Single-cell RNA sequencing analysis of Chenghua pig skin

To characterize the overview single-cell atlas from CH pig skin of different anatomic sites efficiently, we isolated skin cells form six different anatomic sites on the head, ear, shoulder, back, abdomen and leg from three female 180-day-old CH pigs (Figure 1A). After filtering for quality control, we obtained a total of 215,274 cells, which were globally visualized with 21 cell clusters in the t-SNE plot (Figure 1B). On average, 956 genes and 2687 unique molecular identifiers (UMIs) per cell were detected (Figure 1—figure supplement 1A, 1B). The most representative expressed genes for each cluster were used to identify the 21 cell clusters with the heatmap (Figure 1C). The 21 cell clusters constituted seven cell types with known expressed marker genes, of which the SMCs (clusters 0, 2, 5, 6 and 13) were marked by MYH11 and ACTA2, ECs (clusters 3, 4, 7,10 and 11) were marked by PECAM1 and APOA1, FBs (clusters 1, 8, 9 and 12) were expressed by LUM and POSTN, myeloid dendritic cells (MDCs) (clusters 14, 16 and 18) were labeled by BCL2A1 and CXCL8, T cells (TCs) (cluster 15) were highly expressed by RHOH and SAMSN1, KEs (cluster 17) were tabbed by KRT5 and S100A2, and epidermal stem cells (ESCs) (clusters 19 and 20) were stamped by TOP2A and EGFL8 (Figure 1D, 1E and Figure 1—figure supplement 1C).

**Figure 1.**
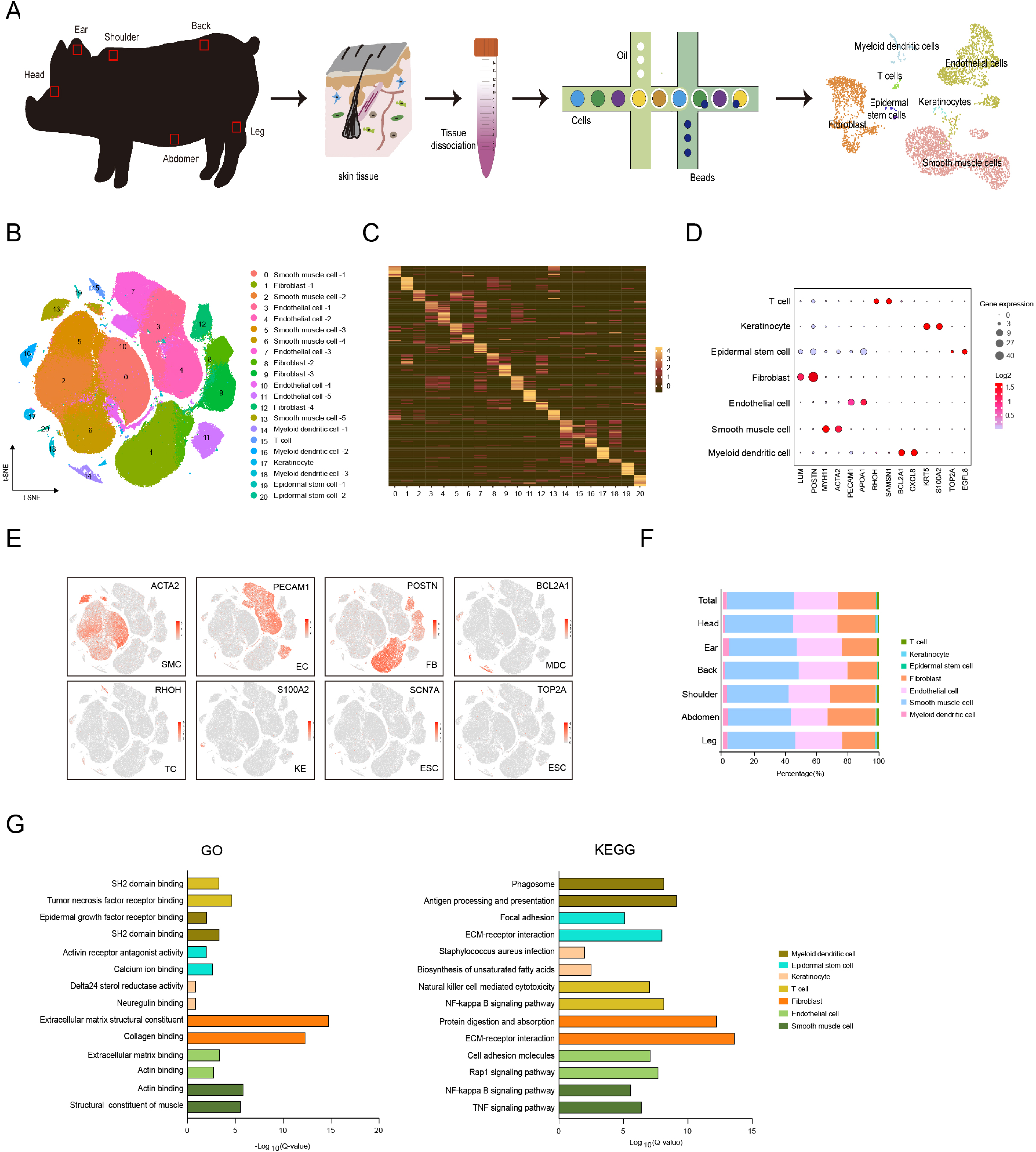
Single-cell atlas of six different anatomical areas of Chenghua pig skin. (**A**) Flowchart overview of skin single-cell RNA sequencing from different anatomical skin regions of Chenghua pig. (**B**) The t-SNE plot visualization showing 21 clusters of annotated cell types from Chenghua pig skin. (**C**) Heatmap showing the top 12 of highly expressed genes from each cluster. Each column represents a cluster, each row represents a gene. Light yellow shows the maximum expression level of genes, and deep green shows no expression. (**D**) Dot plot showing the two representative genes for each cell type. Color indicates the log2 value, and circle size indicates gene expression level. (**E**) The marker genes for each cell type are distributed on the t-SNE plot. Color indicates gene expression. (**F**) The distribution ratio of cell types for total cells and six different anatomical skin areas. (**G**) The most enriched GO terms and KEGG pathways for each cell type. SMC, smooth muscle cell; EC, endothelial cell; FB, fibroblast; MDC, myeloid dendritic cell; TC, T cell; KE, keratinocyte; and ESC, epidermal stem cell. **Figure 1—Source data 1.** Source data of marker genes for each cluster in Figure 1C.

The distribution ratio of these cell types was visualized among total data consisting of 42.9% SMCs, 28.1% ECs, 24.6% FBs, 2.5% MDCs, 0.9% TCs, 0.6% KEs and 0.3% ESCs, with similar distribution trends for the main cell types in various skin regions (Figure 1F). In addition, the cell number and cell identification among the different anatomic skin sites for the head, ear, shoulder, back, abdomen and leg were comparable, which indicated that the cell types displayed subtle differences, but cell number per cell type was significantly varied (Figure 1—figure supplement 2). The marker genes for each cell type showed the dominant transcriptional traits, and the most significantly enriched pathways were presented using Gene Ontology (GO) and Kyoto Encyclopedia of Genes and Genomes (KEGG) analyses (Figure 1G). Significant examples of GO function terms involved in extracellular matrix structural constituent or collagen binding for FBs, actin binding or structural constituent of muscle for SMCs, and extracellular matrix structural constituent or collagen binding to ECs. The pathways are prominently attributed to FBs such as protein digestion and absorption or ECM-receptor interaction, ECs involved in cell adhesion molecules or the Rap1 signaling pathway, and SMCs including the NF-kappa B signaling pathway or the TNF signaling pathway.

Moreover, given the potential cross-species comparisons, we implemented overlapping skin cell atlases among pigs, humans and mice using a t-SNE plot (Figure 1—figure supplement 3A). The captured gene and UMI counts were more advantageous for human skin cells (Figure 1—figure supplement 3B). The cell types were similar for the three species, while the percentage of cell types was different such as smooth muscle cells, endothelial cells or keratinocytes (Figure 1—figure supplement 3A, 3C). Some skin tissue marker genes were showed on the heatmap and dot plots, which examined the shared or species-specific genes in all cell types among the three species (Figure 1—figure supplement 3D, 3E). When discounting the uniqueness of the skin thickness of CH pig breeds resulting in this discrepancy of the cell type proportions such as excessive cell number of SMCs and ECs, dominantly originated from the vessel bed, we believed the pig skin tissue could be considered as the human skin model at skin single cell atlases level for research purposes.

### Heterogeneity of the smooth muscle cells

The most abundant cells were SMCs, followed by the ECs and FBs. SMCs play a critical role in forming blood vessels and arrector pili muscle (APM) of the skin (Driskell et al., 2013; Liu and Gomez, 2019). Previous studies have uncharacterized SMCs in skin tissue. Here, we interrogated the heterogeneity and function of cutaneous SMCs. The t-SNE analysis divided smooth muscle cell into five subpopulations (clusters 0, 2, 5, 6 and 13) (Figure 2A), in which the MYH11 and ACTA2 gene markers were used for the immunohistochemistry staining of skin sections to validate the microanatomical sight of SMCs (Figure 2B). Meanwhile, we performed GO functional analysis using the highly expressed genes of each cluster (Figure 2C). Clusters 0 and 13 predominantly taken part in structural constituent of muscle, acting filament binding and acting binding. The engagement of main inflammatory response and chemokine activity belonged to clusters 2, 5 and 6, of which cluster 2 was also involved in collagen binding and metallopeptidase activity.

**Figure 2.**
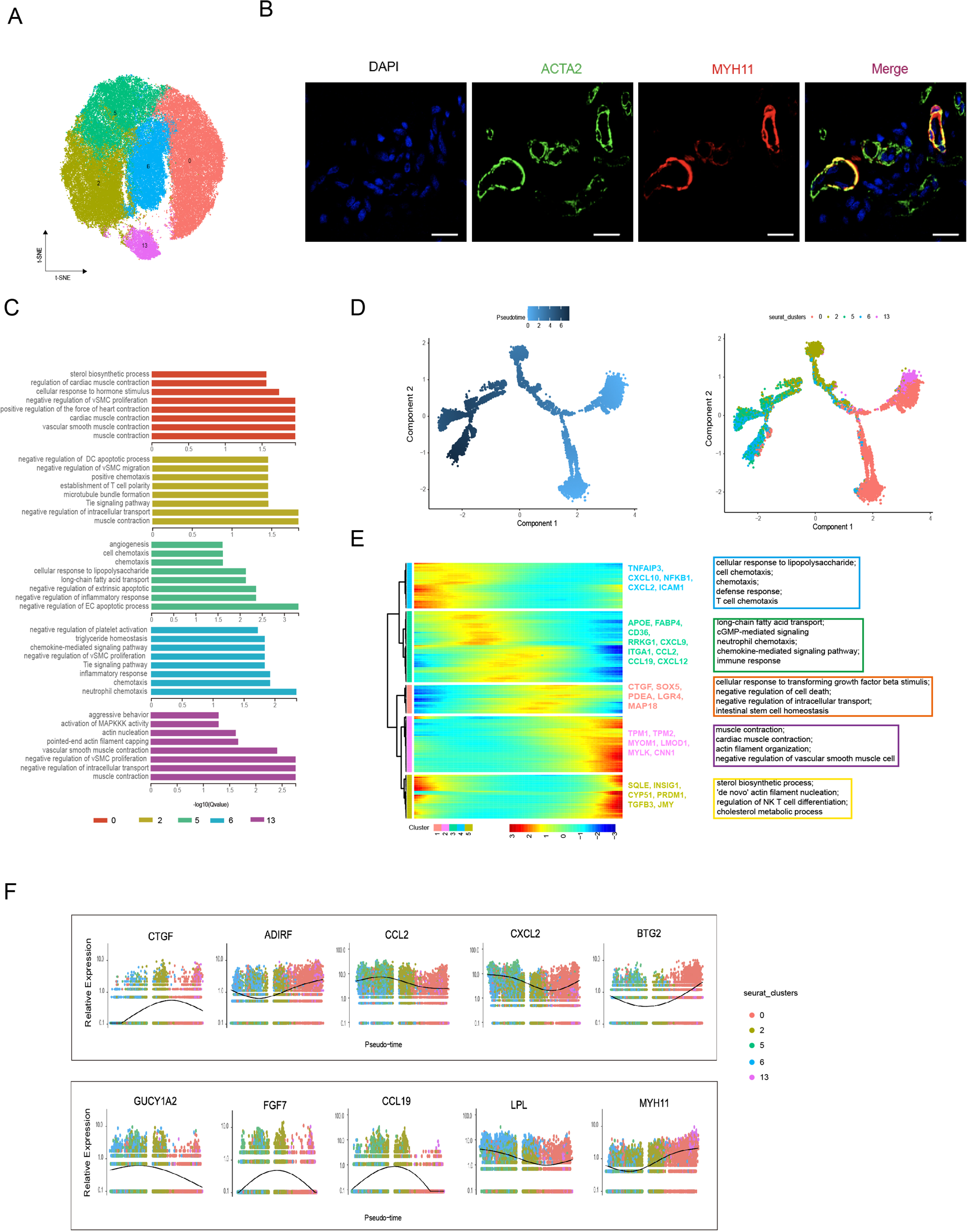
Smooth muscle cell heterogeneity. (**A**) The t-SNE plot visualization of smooth muscle cell populations including clusters 0, 2, 5, 6 and 13. (**B**) Confocal images showing immunofluorescence staining of ACTA2 (green) and MYH11 (red) in back skin sections, representative markers of smooth muscle cells. Scale bar = 50 μm. n = 3. (**C**) The enriched GO terms of biological process for each smooth muscle cell subpopulation sorted by q-value. (**D**) Pseudotime ordering of SMC subpopulations using Monocle 2. (**E**) Heatmap illustrating the dynamics of representative differentially expressed genes among SMC phenotypes, in which the important GO terms relating to biological process were described. (**F**) These genes expression along pseudotiom in SMC subpopulations.

These results showed that SMCs played an important role in blood vessel homeostasis and function, partial collagen binding and immune responses of skin tissue. In previous studies, vascular SMCs displayed a high degree of plasticity and seemed to differentiate into other-like cell types characterized by the expression of marker genes such as mesenchymal-like and fibroblast-like. This evidence, combined with GO function analysis and the expression level of conventional marker genes, such as MYH11 and ACTA2 for SMC, GUCY1A2, CCL19, FGF7 and ASPN for mesenchymal cells (MECs), and LPL, CCL2, IL6 and CXCL2 for macrophages (MACs), presumed that cluster 2 might be mesenchymal-like phenotype or clusters 5 and 6 might represent macrophage phenotype. To further validate the topography of SMC phenotypes, we performed pseudotime trajectory analysis based on the Monocle algorithm (Figure 2D). The trajectory demonstrated that SMCs experienced a dynamic transition from SMCs to mesenchymal-like phenotype and mesenchymal-like phenotype to macrophage-like phenotype. The sequential gene expression dynamics with all branches were visualized and showed five gene sets along expression pattern, which primarily deciphered three cell states (Figure 2E). Gene set 1 and 3 showed high expression of CTGF, LGR4, FABP4, CCL2,CCL19 and FGF7, and enriched GO terms of negative regulation of cell death, intestinal stem cell homeostasis, long-chain fatty and transport and immune response, which conformed well to the mesenchymal-like cells. With high expression of MYH11, MYOM1, TPM1,TPM2, SQLE, BTG2, ADIRF and TGFB3 in GO terms of muscle contraction, actin filament organization and ‘de novo’ action filament nucleation belonged to gene set 2 and 5, which was greatly similar to contractile SMCs. Gene set 4 showed high expression of CXCL10, CXCL2, ICAM1, LPL and IL6, which main were gathered in GO terms of cellular response to lipopolysaccharide, cell chemotaxis and defense response, which may represent macrophage-like cells. Additionally, some cell type-specific marker genes expression trends in five SMCs clusters were presented (Figure 2F), e.g., the MECs-specific genes GUCY1A2, FGF7 and CCL19 were highly expressed in cluster 2, the MACs-specific genes LPL was enriched in clusters 5 and 6, and SMCs-specific genes MYH11 was highly expressed in clusters 0 and 13. These results proved our hypothesis that cluster 2 was mesenchymal-like phenotype or clusters 5 and 6 were macrophage-like phenotype.

For different cutaneous anatomic sites, we found that the total cells number showed a significant difference, while the distribution ratio of smooth muscle cell subpopulations displayed a similar trend, of which the cell number of macrophage-like phenotype was most distinct, followed by SMCs, and that of the mesenchymal-like phenotype was relatively constant (Figure 3A). To decode the transcriptomic changes in SMCs of different cutaneous anatomic sites, the differentially expressed genes (DEGs) were presented among fifteen groups (Figure 3B). The upregulated and downregulated genes of the differentially compared groups were analyzed using GO enrichment terms (Figure 3C). Significant enriched terms of GO analysis terms for upregulated genes primarily referred to extracellular region, collagen-containing extracellular matrix and long-chain fatty acid transport, and downregulated genes took part in cytokine activity, CXCR chemokine receptor binding and positive regulation of T cell migration. The majority of upregulated genes subsisted in back skin compared to other locations, so we implemented KEGG analysis, which involved in PI3K-Akt signaling pathway, MAPK signaling pathway, immune response and integration (Figure 3D). We chose some genes of related ECM-integrins and immune response to present at different skin anatomic sites (Figure 3E), which showed that immune response correlated closely with shoulder skin region or ECM-integrins tightly linked to skin locations on the head, back and shoulder. Moreover, we investigated the key transcription factors (TFs) along all the DEGs among various compared groups due to the importance of gene expression regulators using single-cell regulatory network inference and clustering (SCENIC). The SCENIC algorithm demonstrated a series of main regulons such as EGR1, ATF3, NFKB1, PRDM1 and REL, and related target genes (Figure 3F). TFs, especially ATF3 and EGR1, primarily regulate their target genes at back skin. These results provide well insights into the SMCs heterogeneity in heredity and function in different anatomic sites.

**Figure 3.**
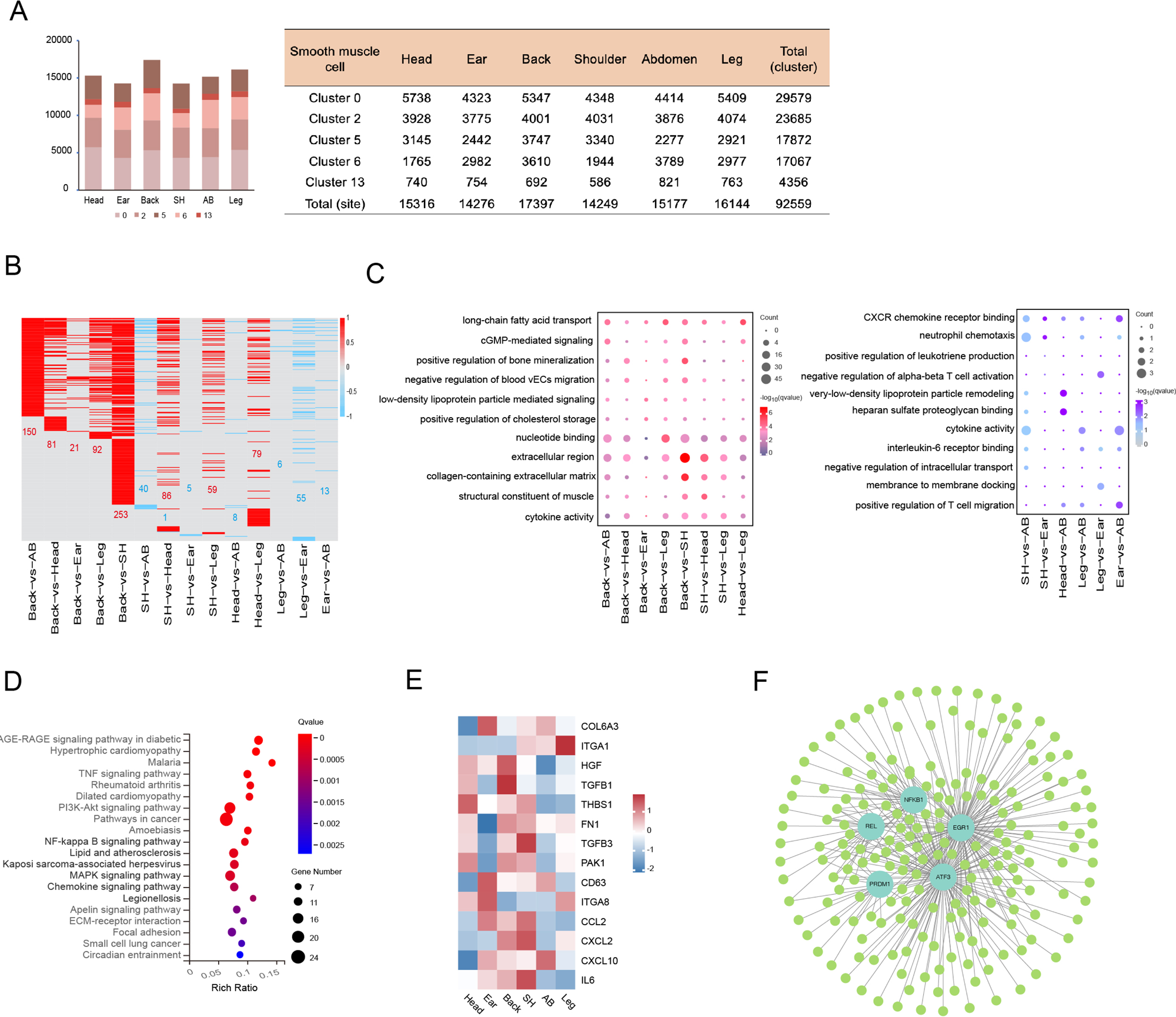
Smooth muscle cell heterogeneity of different anatomical skin regions. (**A**) The cell number of SMC subpopulations in different skin regions. (**B**) Heatmap showing the differentially expressed genes of SMCs in multiple compared groups. Red represents upregulated genes, blue represents downregulated genes and the number of differentially expressed genes is indicated. (**C**) The enriched GO terms of multiple compared groups. Color indicates q-value, circle indicates gene counts. (**D**) KEGG analysis for upregulated genes of back skin compared to other locations. (**E**) The expression level of genes involved in ECM-integrins and immune response pathway in different skin regions. Red represents high expression of genes. (**F**) Transcriptional regulatory network of differentially expressed genes for SMCs in multiple compared groups. Blue nodes represent regulators and green nodes represent the target genes of regulators. **Figure 3—Source data 1.** Source data of the differentially expressed genes of SMCs in multiple compared groups in Figure 3B.

### Heterogeneity of the endothelial cells

The ECs underlying the vascular systems and primarily participate in blood and skin homeostasis (Kalucka et al., 2020). ECs were captured from six different anatomic sites and were classified into five subpopulations in our datasets, which were visualized with the t-SNE plot (Figure 4A). GO functional terms analysis was carried out according to the enriched expression genes of each cluster, which were closely related to some terms of angiogenesis, immune response, response to viruses, cell migration, cell adhesion and regulation of catalytic activity (Figure 4—figure supplement 1A). To validate the spatial position of ECs in dermis, we detected the expressions of representative PECAM1 and APOA1 genes via the immunofluorescence of skin section (Figure 4B). ECs heterogeneity can occur in diverse vascular bed of different anatomic sites or health and disease (Kalucka et al., 2020). Previous studies have reported that endothelial cells (ECs) elaborately construct the vasculature throughout the cutaneous dermis and are classified as arteriole ECs, capillary ECs, venule ECs and lymphatic ECs (Li et al., 2021). Based on the known and reported markers of EC phenotypes (Li et al., 2021; Wang et al., 2022), ECs were composed of arteriole ECs expressing markers SEMA3G and MECOM (clusters 7 and 10), capillaries ECs expressing marker PLVAP (cluster 3), venule ECs expressing markers SELE and ACK1 (cluster 4), and lymphatic ECs expressing markers LYVE1 and PROX1(cluster 11) in dermis (Figure 4C). The pseudotime trajectory analysis of EC phenotypes showed an organized axis of blood ECs starting from arteriole and ending at venule, in agreement with the previous literature (Wang et al., 2022), and pseudotime trajectory also formed an arteriovenous anastomosis tendency (Figure 4D). EC phenotypes exhibit diverse molecule and function such as immune and metabolism trait, as well as appearing in differential tissues and sites, resulting in heterogeneous functions of inter-organ and different sites. Furthermore, we explored the expressions of integrins (focal adhesion), immune (cell adhesion molecules, chemokine signaling pathway, antigen processing and presentation, leukocyte transendothelial migration and Th1 and Th2 cell differentiation) and metabolism (inositol phosphate, mucin type o-glycan biosynthesis, ether lipid, sphingolipid and glycerolipid) across multiple EC phenotypes. ECs related metabolism in our dataset was considerably active in arteriole ECs, especially cluster 10 involving ACER3, which controlled the homeostasis of ceramides, and LCLAT1, a lysocardiolipin acyltransferase regulating activation of mitophagy (Figure 4E). The focal adhesion genes were more significantly upregulated in arteriole ECs and lymphatic ECs compared to other phenotypes, including ACTG1, BIRC3 and THBS1 (Figure 4—figure supplement 1B). In cell adhesion molecules, PTPRM and CDH5, main responsibility for intercellular adhesion between ECs, were highly enriched in arteriole ECs; moreover, PECAM1, SELE and SELP were enriched in venule ECs (Figure 4—figure supplement 1C). Other immune pathways showed that different EC phenotypes significantly high expressed diverse genes, such as CXCL14 (involved in monocyte and recruitment) in capillary ECs, CCL26, CXCL19 and CCL26 in venule ECs (Figure 4—figure supplement 1D). The functional diversity of EC phenotypes showed the degree of ECs heterogeneity.

**Figure 4.**
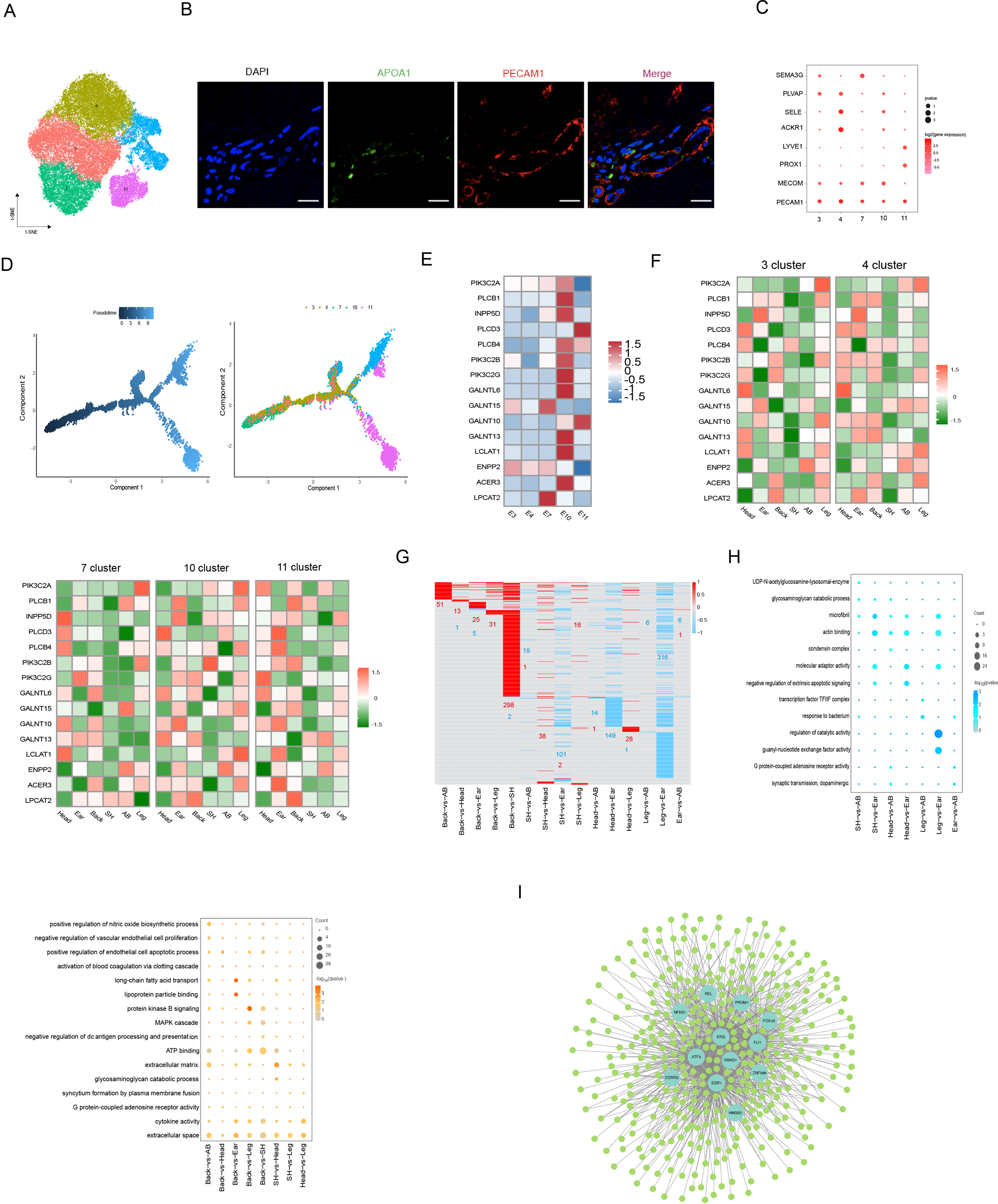
Endothelial cell heterogeneity. (**A**) The t-SNE plot visualization of endothelial cell populations. (**B**) Immunofluorescence staining of APOA1 (green) and PECAM1 (red) in back skin sections, representative markers of endothelial cells. Scale bar = 50 μm. n = 3. (**C**) Dot plot representing marker genes of endothelial cell phenotypes. Color indicates gene expression, circle indicates the log2FC value. (**D**) Pseudotime ordering of ECs subpopulations using monocle 2. (**E**) Heatmap showing the gene expression of metabolic pathways in EC subpopulations. (**F**) Heatmap of gene expression of metabolic pathways in EC subpopulations of different skin regions. (**G**) Heatmap of DEGs for ECs in multiple compared groups. Red represents upregulated genes, blue represents downregulated genes. (**H**) The significantly enriched GO terms of ECs in multiple compared groups. (**I)** Regulatory network of DEGs for ECs of different skin regions. Blue nodes represent regulators and green nodes represent the target genes of regulators. **Figure 4—Source data 1.** Source data of the differentially expressed genes of ECs in multiple compared groups in Figure 4G.

Here, we found that the cell number of EC phenotypes was difference among different anatomic sites, with the back skin holding the most arteriole ECs and minimal lymphatic ECs (Figure 4—figure supplement 1E, 1F). To further confirm the heterogeneity of EC phenotypes in the skin sites of head, ear, shoulder, back, abdomen and leg, we compared these genes expression of integrins, immune and metabolism pathways (Figure4 F and Figure 4—figure supplement 2A-C). For example, for the metabolism pathway, compared to other sites, the activity of capillaries ECs, venule ECs and arteriole ECs (7 cluster not including 10 cluster) was depressed in shoulder skin, while high activity in capillary ECs, venule ECs and arteriole ECs was showed in leg skin including ACER3 and PIK3C2A enhanced cell viability (Gulluni et al., 2021), and high activity in lymphatic ECs, arteriole ECs (10 cluster not including 7 cluster) and venule ECs was presented in ear skin including ACER3, GALNT10 and PIK3C2B, a member of class II PI3Ks controlling cellular proliferation, survival and migration. With abundant results on the related pathways expression for EC phenotypes in different sites showed the heterogeneity of ECs for different anatomic sites of skin.

To uncover the underlying molecular mechanisms driving the differential skin sites of ECs, we compared the DEGs with differentially compared groups among different anatomic sites (Figure 4G) and GO terms analysis was implemented for upregulated and downregulated genes (Figure 4H). Enriched terms relating to long-chain fatty acid transport, lipoprotein particle binding and extracellular matrix were showed in upregulated differential genes, while downregulated differential genes main were existed in terms involving regulation of catalytic activity, acting binding and molecular adaptor activity. Of note, the CD36, a multifunctional fatty acid transporter, was reported related metabolic state of fibroblasts for ECM regulation (Zhao et al., 2019). Here, we found that CD36 was upregulated in the back compared with others except head, which was enriched in metabolic terms such as long-chain fatty acid transport and regulation of nitric oxide. FABP4, fatty acid-binding protein 4, was a lipid transport protein that was significantly differentially expressed in nine pairs compared groups. Pioneering study showed FABP4 was strongly expressed in subcutaneous adipocytes and adipose ECs (Wang et al., 2022). Combining data showed skin thickness might have a positive correlation with subcutaneous fat deposits. Additionally, we constructed single-cell transcription-factor regulatory network with all DEGs (Figure 4I). The analysis predicted the following main transcriptional factors: ATF3, EGR1, ERG, FLI1, PRDM1, and NFKB1. The regulation of ATF3, EGR1 and ERG TFs predominate were exited in compared groups of back vs. shoulder and leg vs. ear skin. With these finding, we presented the heterogeneity of ECs in different anatomic sites of skin.

### Heterogeneity of the fibroblast

The dermal fibroblasts synthesize the ECM that forms the connective tissue of skin dermis to maintain the skin morphology and homeostasis. We found the CH skin thickness of differential skin sites owed striking difference such as back skin thickness on average at 5.48 mm and that ear at 1.52 mm (Figure 5—figure supplement 1A). In terms of overall skin section, the skin histomorphology of different anatomic sites exhibited some difference in sparsity of collagen fibers or the number of appendages, and dermal thickness descended from the back, head, shoulder, leg, abdomen to ear (Figure 5A and Figure 5—figure supplement 1B). Curiously, we inquired whether the discrepancy in ECM accumulation in different skin sites was caused by fibroblast heterogeneity. Next, fibroblasts maps were presented from six different skin anatomic sites using the t-SNE plot, which was established by four clusters (clusters 1, 8, 9 and 12) (Figure 5B), and the cell number of clusters was estimated (Figure 5C and Figure 5—figure supplement 1C). In previous reports (Philippeos et al., 2018; Solé-Boldo et al., 2020), fibroblast in cluster 1 highly expressed MGP and MFAP5, known markers of reticular fibroblast, the most representative markers of COL6A5, WIF1 and APCDD1 of papillary fibroblast in clusters 8 and 9, and the mesenchymal subpopulation signature was typically characterized by enriched expressed CRABP1, TNN and SFRP1 in cluster 12. GO analysis showed the functions were closely related with extracellular matrix organization, collagen fibril organization and cell adhesion, which illustrated four clusters of fibroblasts owned analogous functions in our dataset (Figure 5—figure supplement 1D). Likewise, the label-LUM and POSTN genes were marked on fibroblasts of skin section via immunofluorescence (Figure 5D).

**Figure 5.**
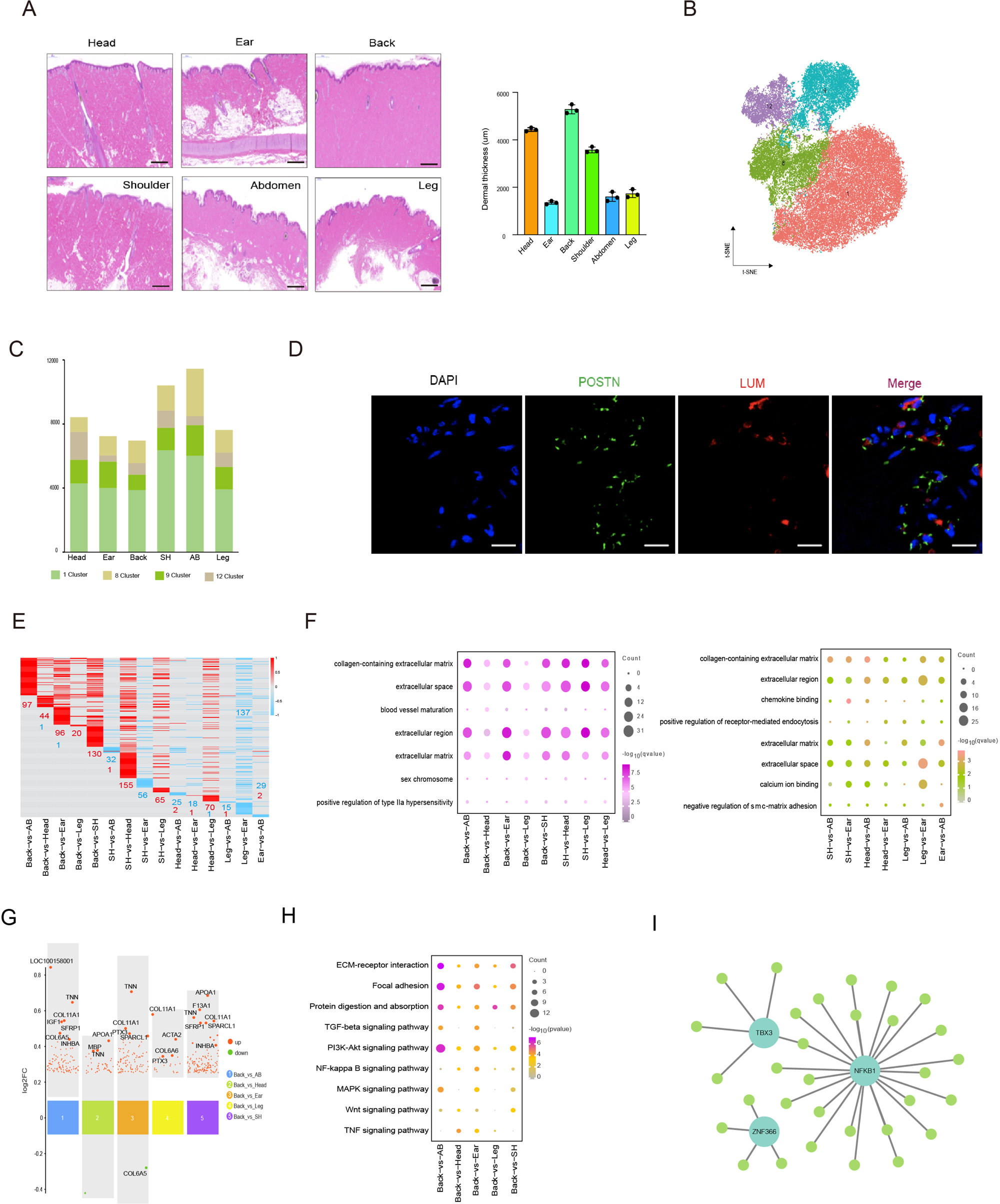
Fibroblast heterogeneity. (**A**) Skin section with HE staining (left) and dermal thickness of six different sites (right) including head, ear, back, shoulder, abdomen and leg. Scale bar = 1000 μm. n = 3. (**B**) The t-SNE plot showing FB populations. (**C**) The cell number of FB populations in different skin regions. (**D**) Images showing immunofluorescence staining of POSTN (green) and LUM (red) in back skin sections, representative markers of FBs. Scale bar = 50 μm. n = 3. (**E**) Heatmap of DEGs for FBs in multiple compared groups. Red represents upregulated genes, blue represents downregulated genes. (**F**) The enriched GO term of FBs in multiple compared groups. (**G**) Multiple volcanic maps showing the DEGs of compared groups in back skin compared to other locations. Representative genes are indicated. (**H**) KEGG analysis of representative genes in image G. (**I**) Regulatory network of DEGs of FBs of different skin regions. Blue nodes represent regulators and green nodes represent the target genes of regulators. **Figure 5—Source data 1.** Source data of the differentially expressed genes of FBs in multiple compared groups in Figure 5E.

With the discrepancy in ECM accumulation in different skin sites, we excavated the upregulated and downregulated DEGs among the diverse compared groups by heatmap, in which all DEGs were upregulated in back skin compared with other skin sites (Figure 5E). The remarkable GO enrichment terms of extracellular matrix, extracellular region, extracellular space and collagen-containing extracellular matrix showed all compared groups using GO function analysis, suggesting the difference in gene expression level of FBs resulted entirely in extracellular various in different skin sites (Figure 5F). We further explored the key gene causing the discrepancy in ECM accumulation, so the top DEGs were visualized in the compared groups (Figure 5G and Figure 5—figure supplement 1E). The point photograph presented some overlapping genes in multiple compared groups especially back skin compared with other skin sites, including TNN, COL11A1, SFRP1, COL6A5, INHBA, APOA1, IGF1 and SPARCL1. Notably, TNN, called tenascin-N(W), is lager domain glycoprotein that has the potential to modify cell adhesion and typically contribute to cell motility (Chiquet-Ehrismann and Tucker, 2011); COL11A1, an extracellular matrix structural constituent, comprises a subclass of regulatory collagens fibrillogenesis that synergistically assemble other types of collagen such as collagen I, determining fibril structure, fibril organization and functional traits (Smith and Birk, 2012; Sun et al., 2020); SFRP1, a member of secretory glycoprotein SFRP family, is regarded as one of the main classes of macromolecules making up the ECM elements and is reported to be an antagonist that inhibits human hair follicles recession (Bertolini et al., 2021; Jiang et al., 2022); INHBA is a member of TGFβ superfamily and is modified by AP1 expression (Ham et al., 2021). Subsequently, we implemented KEGG analysis in the compared groups (Figure 5—figure supplement 1F, G) and presented dominant enrichment pathways such as ECM-receptor interaction, focal adhesion, protein digestion and adsorption and TGF-beta signaling pathway. Interestingly, these typical overlapping genes were tightly connected with ECM production (Figure 5H). At a consequence, TNN, COL11A1 and INHBA were considered key candidate genes for provoking ECM accumulation. In addition, The SCENIC algorithm demonstrated NF-kB1, TBX3 and ZNF366 regulons regulated some DEGs in FB population (Figure 5I). Of note, the targeted INHBA is targeted by TBX3 regulons. Together, our results showed the heterogeneity of FBs in different anatomic sites of skin.

### The difference of back skin cells between Chenghua and Large White pig

In our pervious report, the skin of CH pig was thicker than that of Large White (LW) pigs (8.5 mm vs. 3.0 mm) (Zou et al., 2022), which was consistent with the current data (Figure 6A and Figure 6—figure supplement 1A). Curiously, the pattern of heterogeneity in skin cell for different skin anatomic sites is whether also exist in different breed. To further verify the difference in skin cells across breeds, we also implemented single-cell sequencing for the back skin of LW pig. We received a total 18,441 cells after removing minimum count cells, doublet cells and more than 5% cell-contained mitochondrial genes. The t-SEN analysis revealed the 18 clusters composed of six cell types including SMCs (clusters 1, 2, 5, 6, 9, 10 and 14), ECs (clusters 0, 3, 4, 12 and 15), FBs (clusters 7, 8, 11 and 13), lymphatic cells (LYCs) (cluster 16), Langerhans cells (LHCs) (cluster 17) and ESCs (cluster 18) between CH and LW pig (Figure 6B), and the marker genes of cluster were showed in Figure. 6C. The genes/UMIs per cell and distribution of cell types were compared between the two breeds (Figure 6D and Figure 6—figure supplement 1B-D). The main cell types were still SMCs, ECs and FBs, and we compared the DEGs in two breeds (Figure 6E, F), which showed a large difference was in FB populations. KEGG analysis for the main three cell types manifested significant pathway of ether lipid metabolism for ECs including LPCAT2 and ENPP2 genes, PPAR signaling pathway for SMCs, and PI3-Akt signaling pathway, protein digestion and absorption, ECM-receptor interaction, focal adhesion and TGF-beta signaling pathway for FBs involving in TNN, POSTN, COL11A1, IGF1 and INHBA genes that overlapped with DEGs of FBs of CH pig skin (Figure 6 G and Figure 6—figure supplement 1E). Moreover, the extracellular space and extracellular region part were the representative striking terms for DEG of FB population by GO terms analysis (Figure 6H). An analysis of the data proved the ECM accumulation in skin tissue was probably dependent on these overlapping genes, which might be not bound up with the origin of anatomical regions or breed.

**Figure 6.**
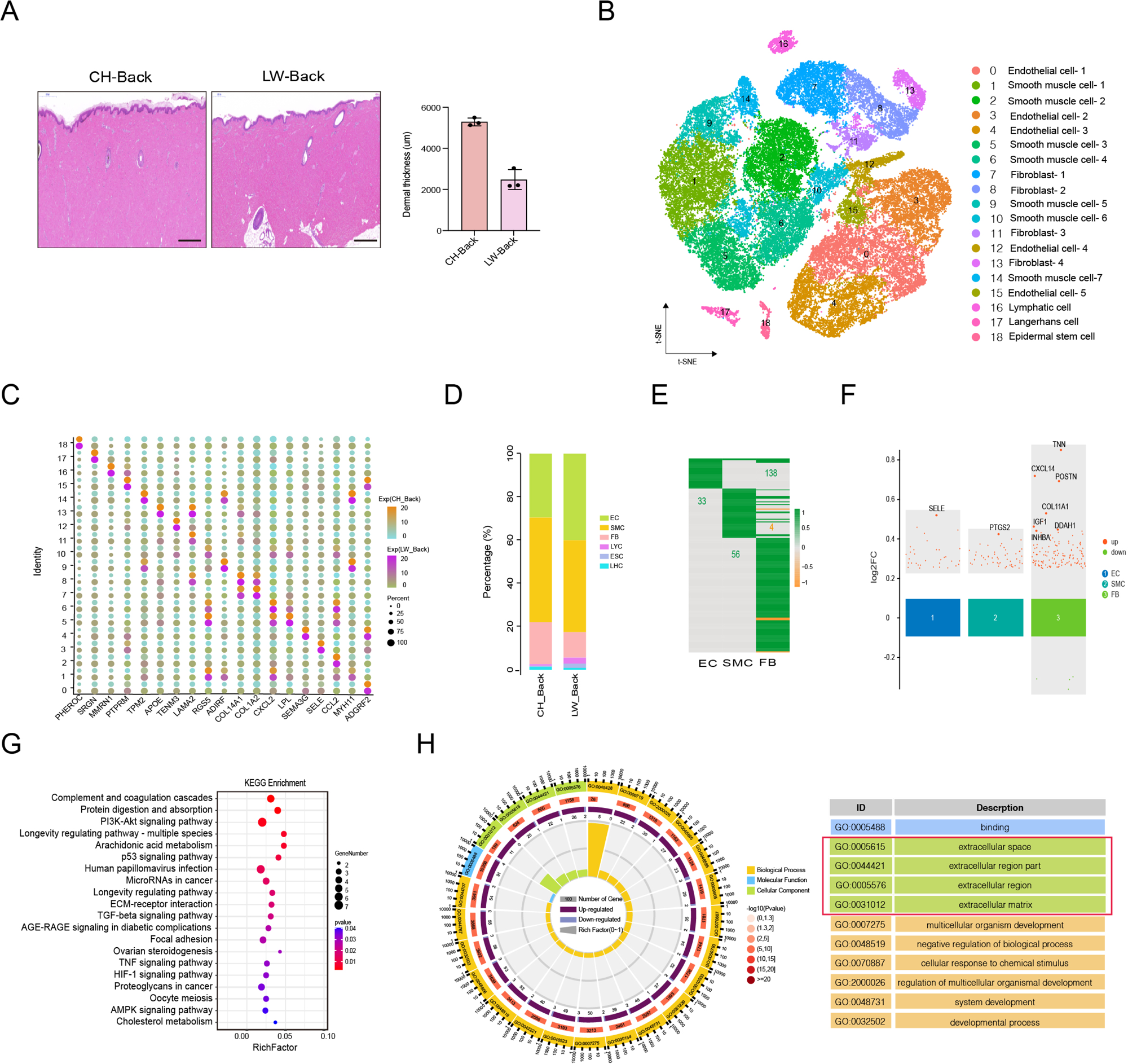
The difference in back skin cells between Chenghua and Large White pigs. (**A**) Skin section with HE staining (left) and dermal thickness of back skin between CH and LW pig (right). Scale bar = 1000 μm. n = 3. (**B**) The t-SNE plot visualization of all clusters of annotated cell types between CH and LW pigs. (**C**) Representative genes of each cluster of skin cells between CH and LW pigs. Color represents the gene expression and circle represents the percentage of cells. (**D**) The distribution of cell types between CH and LW pig skin tissues. (**E**) Heatmap of DEGs for SMCs, ECs and FBs. Green represents upregulated genes, orange represents downregulated genes. (**F**) Multiple volcano maps of DEGs for SMCs, ECs and FBs. Representative genes are indicated. (**G**) KEGG analysis of DEGs for FBs. (**H)** GO term of DEGs for FBs. Red region is the most enriched GO terms. **Figure 6—Source data 1.** Source data of marker genes for each cluster in Figure 6C. **Figure 6—Source data 2.** Source data of the differentially expressed genes of SMCs, ECs and FBs in compared group in Figure 6E.

### The communication of overview skin cell

Intercellular communication plays an important role in complex tissues. Understanding cell-cell communication in skin tissue requires accurate signaling crosstalk via ligands, receptors and their cofactors, and effective overview analysis of these signaling links. To investigate the signaling crosstalk of seven cell types in skin tissue, we established intercellular communication by the R package CellChat. The seven cell types were deemed as communication “hub”, which detected 547 ligand-receptor pairs and were further categorized into 36 signaling pathways including the COLLAGEN, LAMININ, FN1, PDGF, CCL, CXCL, MIF and ITGB2 pathways (Figure 7A). Specifically, the COLLAGEN and LAMININ pathway exhibited highly abundant signaling interactions among seven cell types. Network centrality analysis of the COLLAGEN/LAMININ pathway revealed FB populations were the main source of the COLLAGEN/LAMININ ligands targeting SMC and ESC populations, which showed the COLLAGEN/LAMININ interactions were primarily paracrine (Figure 7B, C and Figure 7—figure supplement 1A). Importantly, these results reported the elaborately relevance between FBs and SMCs with majority ligand of COL1A1and COL1A2, receptor of CD44 and ITGA1+ITGB1 in the COLLAGEN pathway (Figure 7—figure supplement 1B). Likewise, the LAMININ pathway also showed an analogous phenomenon between FB and SMC populations via ligands LAMC1and LAMB1, which were receptors of CD44 and ITGA1+ITGB1 (Figure 7—figure supplement 1B).

**Figure 7.**
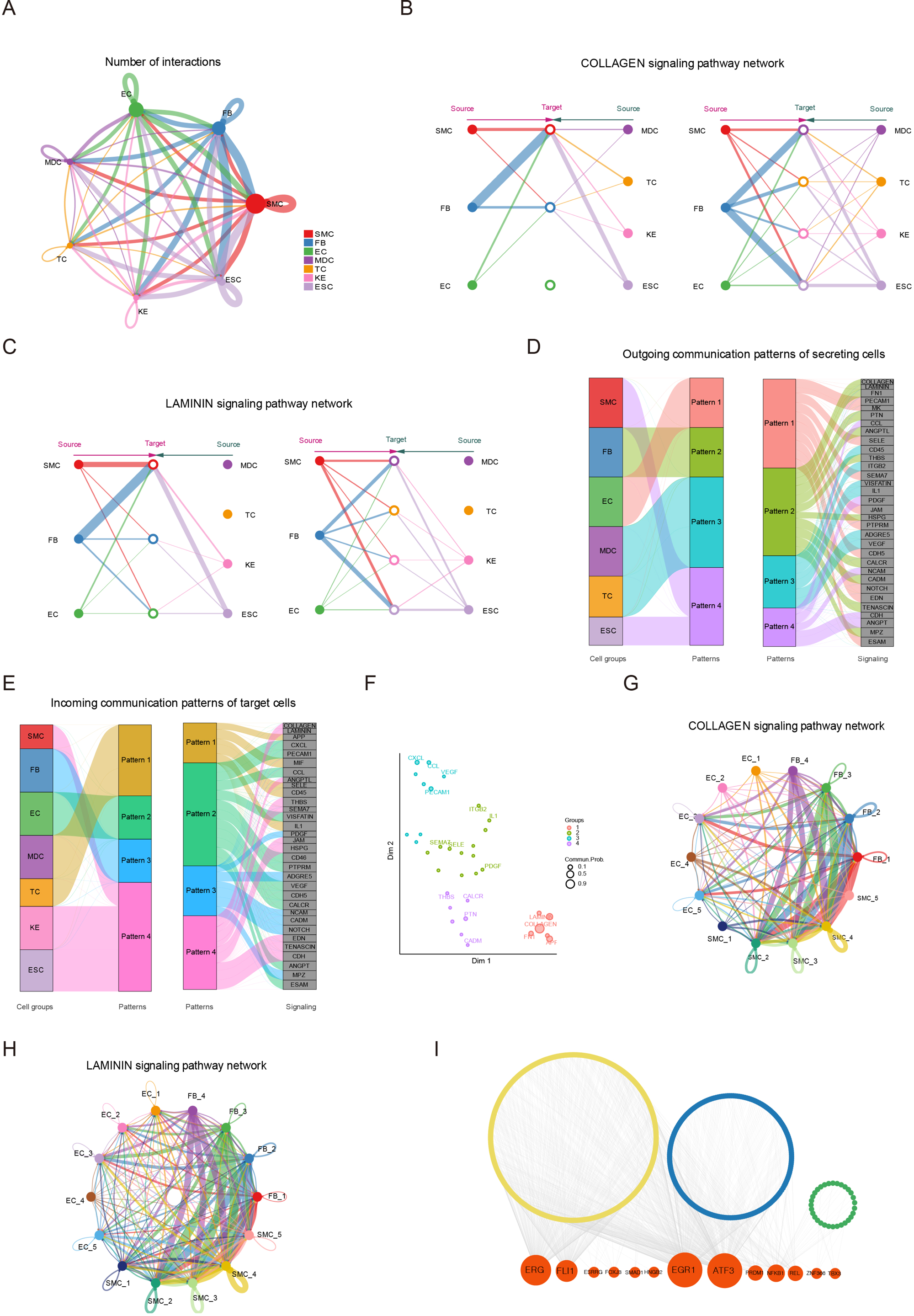
The communication of skin cells. (**A**) Circle plot representing the cell communication among cell types. Circle sizes represent the number of cells and edge width represents the communication probability. Hierarchical plot showing the intercellular communication network for the COLLAGEN (**B**) /LAMININ (**C**) signaling pathways. Circle sizes represent the number of cells and edge width represents communication probability. The inferred outgoing communication patterns (**D**) and incoming communication patterns (**E**) of secreting cell of CH pig skin. (**F**) The distribution of signaling pathways with their functional similarity. The COLLAGEN (**G**) /LAMININ (**H**) signaling network among cell subpopulations of SMCs, FBs and ECs. Circle sizes represent the number of cells and edge width represents communication probability. (**I**) Regulatory network of DEGs for SMCs, ECs and FBs of different skin regions. Orange nodes represent regulators and yellow/blue/green nodes represent the target genes of regulators. **Figure 7—Source data 1.** Cell communication of skin cells in Figure 7A.

We implemented a communication pattern analysis to uncover the four patters in outgoing secreting cells or incoming target cells (Figure 7D, E). Outgoing FB populations signaling was identified by pattern #2, which represented multiple pathways such as COLLAGEN, LAMININ, FN1, PTN, ANGPTL and THBS. Outgoing SMCs and ESCs signaling was characterized by pattern #4, included in CDH, ANGPT and PDGF pathways. Outgoing ECs signaling was characterized by pattern #1, which was involved in PECAM1, MK and NOTCH pathways. The pattern #3 presented CD45, IL1 and VEGF pathways for outgoing MDC and TC populations signaling. For incoming communication target cells pattern, incoming FB populations signaling was characterized by pattern #3, representing NCAM, CADM and MPZ pathways. The incoming SMCs, KEs and ESCs signaling was characterized by pattern #4, and that of ECs was characterized by pattern #2.

Furthermore, the signaling pathways were grouped according to their similarity in function or structure. The functional similarity grouping was classified into four groups (Figure 7F). Group #1 and #4, which dominating included COLLAGEN, LAMININ and PTN pathways, largely showed signaling from FBs to SMCs and ESCs. Group #3 dominantly drove PECAM1, CXCL and CCL pathways, which represented the acquisition signaling pathway of ECs. The structural similarity grouping also was identified four groups (Figure 7—figure supplement 1C). To further elaborately explore the communication among FB, SMC and EC subpopulations, we analyzed the COLLAGEN/ LAMININ pathway in the three populations including 14 clusters (Figure 7G, H and Figure 7—figure supplement 1D, E). The network centrality analysis showed clusters 2, 5 and 6 of SMCs and cluster 7 of ECs likely actively take part in cell communication via the ligand of COL1A1/LAMA2, receptor of ITGA1+ITGB1 in the COLLAGEN/ LAMININ pathway. Moreover, the SCENIC algorithm demonstrated NFKB1 as the common regulon among DEGs of three cell types, and the EGR1 and ATF3 regulons regulated the target genes in SMCs and ECs (Figure 7I). These results manifested the communication of skin cells especially main three cell types of SMCs, ECs and FBs.

## Discussion

With the development of high-resolution single-cell sequencing that is applied to delineate the atlas of diverse cell type populations and determine the molecular basis underlying the heterogeneity in many complicated biological processes associated with physiology and pathology among species, especially humans (Han et al., 2020), mice (Kalucka et al., 2020), monkeys (Han et al., 2022) or pigs (Wang et al., 2022). The origin of tissue, its development state or anatomical structure are conductive to heterogeneity of cells. Previous study uncovered the skin scRNA-seq datasets from embryonic development (Ge et al., 2020), different age stages (Zou et al., 2021) and wound healing stages (Guerrero-Juarez et al., 2019) in humans or mice. However, the single cell transcriptional diversity of different anatomic skin regions has not been understood and is caused by different origins of the body skin dermis. Pig as an animal model of human medicine that demonstrate promising alternative cutaneous organ based on their similar physiology, anatomic structure and genetics with humans (Perleberg et al., 2018).

Therefore, in our study, using scRNA-seq detailed analyses of transcriptional similarity, we depict a detailed single-cell atlas of pig skin cells from six different anatomic skin sites involving in head, ear, shoulder, back, abdomen and leg. Compared with reported skin cell types of pig (Han et al., 2022), the cell types varied slightly, but the distribution ratio of cell types was significant different such as 7.2% SMCs, 6.5% ECs, and 45.5% FBs in the reported literature and 42.9% SMCs, 28.1% ECs, 24.6% FBs in this study. Similarly, melanocytes, Schwann cells, mast cell and neural cell were not identified in our datasets, while they were identified in human or mouse skin samples. By the way, SMCs and pericytes, called mural cells in vessels, were unable to precise discriminate between the two cell types because of confusable hallmarks and functions in skin tissues, so we only identified SMCs in our data. Here, we believe that the discrepancy in captured cell types and cell type proportions is based on the cutaneous thickness trait of CH pig breed and different scRNA-seq platforms captures. In this study, we analyzed main three cell types: SMCs, ECs and FBs.

SMCs constitute blood vessels and APM in skin tissue, with a greater proportion in blood vessels. Here, five SMC subpopulations are verified and then separate into three SMC phenotypes. Sophisticated studies have shown SMC phenotypic switching under pathological processes or injured conditions, a way in which SMC shift between contractile phenotype and other type cell phenotypes such as mesenchymal-like, fibroblast-like, macrophage-like, adipocyte-like and osteogenic-like (Yap et al., 2021; Yu et al., 2022). Subsequently several studies found multiple SMC phenotypes including Scal-positive vascular SMC-lineage also existed in healthy vessels (Dobnikar et al., 2018). Interestingly, SMC phenotypic switching occurred in pig skin tissue, varying from contractile SMCs to mesenchymal-like, mesenchymal-like to macrophage-like, with the expression level of marker genes for cell types and function analysis. During physiologic and pathological angiogenesis, macrophages are regarded as a facilitator of vascular integrity and derivatives by way of cytokine secretion and ECM remodeling (Barnett et al., 2016; Debels et al., 2013), which implies SMCs are deemed immune system’s line of defense and positively participate in the immune response in skin tissue. From different skin regions, the cell number of the macrophage-like phenotype is highest at back site, followed by the abdomen, but immune related-genes are primarily existed in shoulder, back and ear, which shows activity of the macrophage-like phenotype might depend on intrinsic factors as well as environmental factors.

Depending on the properties of diverse molecular and function, such as immune responses and metabolic process in ECs, ECs heterogeneous characteristics have been investigated in some organs of human skin (Li et al., 2021), the mouse brain (Kalucka et al., 2020), and pig adipose tissue (Wang et al., 2022), of which ECs diversity remain largely unrevealed in skin tissue from different anatomical locations. In our cutaneous datasets, five subpopulations are identified with ECs and divide into four EC phenotypes, which are placed in order at the pseudotime trajectory indicating the distributed paths of blood vessel, such as arteriovenous anastomosis with vein. Here, an additional level of heterogeneity was explored when analyzing the expression level of pathway genes involved in integrins, immune and metabolism in EC subtypes as well as the EC subtypes in different anatomical regions. In integrins, ITGA6 is high expressed in pig dermis ECs, in accordance with human dermis ECs (Li et al., 2021), and the expression of ITGA6 is significant added in venule ECs and arteriole ECs (cluster 10 not cluster 7) of the pig ear skin site and in capillaries ECs of humans. Cell adhesion molecules are compared to find capillaries ECs and arteriole ECs (cluster 7 not cluster 10) of the back and shoulder skin main enriched MHC class II genes such as SLA-DQB1 and SLA-DRB1, which were highly expressed in lung organ of humans/mice, indicating a role in immune surveillance (Goveia et al., 2020). A funny question regarding the high expression of MHC class II genes in term of slight tissue rejection by blocking MHC class II on human endothelium (Abrahimi et al., 2016) is whether there is a preference for skin graft from specific skin regions, to transplant pig skin into humans. SELE, SELP and ICAM1 main mediate the communication between leukocyte and ECs and are high expressed in venule ECs and arteriole ECs of head, ear and back skin of pig, while the three genes are primarily existed in post-capillary venule ECs of human dermis (Li et al., 2021). CDH5, an intercellular tight junction protein in ECs, is high expressed in arteriole ECs of the ear skin of pig, in keeping with human dermal EC phenotype. Other immune response representative genes such as cytokines (CXCL14, CCL26, CCL24, CXCL12 and CXCL19) that participate in immunocyte recruitment (e.g., neutrophils) or are responsible for the host defense against viral infection, enhancing immune progression and metastasis (Fajgenbaum and June, 2020; Wu et al., 2020), of which CCL24/CCL26, the role of eotaxins (Provost et al., 2013), are enriched in venule ECs and major distributed in the head/abdomen skin regions respectively. For the ECs metabolism pathway, most metabolic genes are significant expressed in arteriole ECs and exhibited overlapping and specific among different skin sites. Interestingly, ENPP2 of lipid metabolism gene reported enhanced the cytokine production (Grzes et al., 2021) and was overexpressed during chronic inflammatory (Argaud et al., 2019), and it was enriched in the abdomen skin, while LPCAT2 of the other lipid metabolism gene under study was positively correlated with lipid droplet content in colorectal cancer (Cotte et al., 2018), which was main highly expressed in back and shoulder skin. These finding demonstrated the extensive phenotypic plasticity and gene expression signatures of all kinds of pathways in different skin sites.

FBs are mesenchymal cells that synthesize ECM of connective tissues, which are responsible for structural integrity, wound repair and fibrosis in skin. Providing plentiful proofs showed FBs heterogeneity is involved in diverse subpopulations such as papillary FBs, reticular FBs, mesenchymal FBs and pro-inflammatory FBs, and its functions in humans and mice (Guerrero-Juarez et al., 2019; Zou et al., 2021). Through the known marker of FB subpopulations, we found three subpopulations not including pro-inflammatory in our dataset, guessing the immune function of SMCs or ECs might replace pro-inflammation FBs due to the enormous cell number of SMCs or ECs. Previous showed FBs of distinct anatomic locations exhibited detectable differences in metabolic activity (Castor et al., 1962) and genome-wide gene expression profiling of 43 skin sites (Rinn et al., 2006). The fact is that three FB subpopulations focused on extracellular matrix organization and collagen fibril organization, resulting in the discrepancy in ECM deposition in different anatomical skin sites.

Therefore, the overlapping remarkable upregulate-genes were found among multi compared groups, especially the back when compared with other areas, and they might be regarded as key genes in ECM deposition. The ECM protein TNN is high expressed in dense connective tissue such as cartilage, adult skeleton and bone (Chiquet-Ehrismann and Tucker, 2011). TNN distinctly located with collagen 3 fibers plays a crucial role in periodontal remodeling, an example of a dense scar-like connective tissue enriched the nerve fibres replacing alveolar bone around the incisor by deficient TNN in mice (Imhof et al., 2020). Here, TNN took part in these pathways that were closely related with skin dermis such as ECM-receptor interaction and PIK-Akt signaling pathway and were significantly upregulated in multiple compared groups uniformly, surmising TNN might a key candidate gene. Collagen I is the most abundant structural macromolecule in skin tissue, and collagen mechanism is determined by minor component as a regulator (Hansen and Bruckner, 2003). Collagens I and XI can package into composite fibrils by nucleation and propagation, in which the collagen XI content is closely connected with collagen I, determining its organization and function properties. Collagen XI is the main factors in collagen I-containing tissue including tendons and cartilage, but not skin tissue, and the absence of COL11A1 expression results in the disruption of fibril phenotype for mature tendons (Blaschke et al., 2000; Sun et al., 2020). INHBA play an important role in the TGF-beta signaling pathway, stimulating the activity of SMAD2/3 and encouraging cell proliferation and ECM production. INHBA expression was significantly upregulated in keloid FBs compared to normal dermal FBs (Ham et al., 2021). Interestingly, overlapping genes including TNN, COL11A1, SFRP1, INHBA were pronouncedly expressed in mesenchymal FBs. With the paradigm of human skin case presented a series of genes were significantly increased in keloid mesenchymal FBs in contrast to normal scar, such as COL11A1, SFRP1, TNC, INHBA, FN1, IGF1, THBS4 and POSTN, suggesting these genes might promote ECM production (Deng et al., 2021). Likewise, these TNN, POSTN, COL11A1, IGF1 and INHBA genes were significantly upregulated in back skin of CH pig compared with back skin of LW pig. Although the mechanisms of physiological skin thickness, fibrosis or scarring (pathological chronic inflammatory) are not all the same, excess ECM accumulation occurs, indicating individual and mutual genes. Therefore, in our study, we speculate TNN, COL11A1 and INHBA expression might play a critical role for the morphology and quantity of collagen fibril-stimulated ECM deposition in skin tissue.

Skin physiological and pathological (wound healing or fibrosis) conditions not only determine the complex and diverse cellular composition but also establish the central signaling pathways between interacting cell groups, offering good insights into cellular crosstalk. For mouse skin wound tissue, network analysis categorized into 25 signaling pathways involving in TGFβ, non-canonical WNT, TNF, SPP1 and CXCL, and identified the inferred TGFβ signaling as the most prominent pathway between myeloid cells and FB populations (Guerrero-Juarez et al., 2019). Twenty-two signaling pathways of embryonic mouse skin were identified, such as WNT, ncWNT, TGFβ, PDGF, NGF, FGF and SEMA3, predicting the WNT signaling pathway paid an important role between epidermal to dermal cells to form skin morphogenesis (Gupta et al., 2019). Moreover, the major highly active pathways in diseased human skin including MIF, CXCL, GALECTIN, FGF and CCL, which showed MIF signaling pathway was main pathway from inflammatory FBs to inflammatory TCs (He et al., 2020). In our datasets, 36 signaling pathways were presented involving in COLLAGEN, LAMININ, FN1, PDGF, CCL, CXCL and MIF, of which the COLLAGEN and LAMININ signaling were the most enriched among different skin regions of pig. These results indicate the key signaling pathways depended on skin morphogenesis.

In summary, in our study, the heterogeneity of main cell types from different anatomic skin sites was comprehensive detailed, giving clear evidence of the use of pig as an excellent skin model focused on generation, transmission, positional information and transplant, paving the foundation for skin priming.

## Materials and Methods

### Skin samples dissociation and cell collection

Skin samples were obtained from three CH pigs at six different anatomical body areas (head, ear, shoulder, back, abdomen, leg) and three LW pigs with one region (back). The fresh skin samples were thoroughly scraped off the hair and subcutaneous fat and were washed thrice with ice-cold Dulbecco’s Phosphate-Buffered Salline (1×DPBS). The skin samples (size approximately 2 cm×2 cm) were fully dissected into small pieces in 4 mL tube and then transferred into 50 mL centrifuge tube with 15 mL mix digestion medium containing 1 mg/mL collagenase type I, II, IV, V (Sigma-Aldrich, Saint Louis, USA), 1 mg/mL elastinase (Coolaber, Beijing, China), and 2 U/mL DNase I (Coolaber, Beijing, China) in Dulbecco’s Modified Eagle Medium (DMED). The skin samples were digested at 37 °C for 120 min-180 min, and simultaneously gently shaken once every 10 min. The digestion reaction was interrupted by DMEM including 10% fetal bovine serum (FBS) (Gibco, New York, USA). Then, the tissue suspension was filtered with 70 μm and 40 μm cell strainer and transfected into a 15 mL centrifuge tube to obtain cells sediment by centrifugation at 350×g for 5 min at 4 °C. The cells sediment was added to 2 mL Red Blood Cells Lysis Solution (Qiagen, Duesseldorf, Germany) at room temperature for 5 min to remove red blood cells. The cells sediment was added to 2 mL TrypLE (Gibco, New York, USA) at 37 °C for 45 min to dissolve cell clot. The dissociated cells were washed twice and resuspended in cold DMED supplemented with 10% FBS. Finally, cells staining with 0.4 % Trypan Blue Solution was used to estimate cell activity rate and concentration by Countess™ Cell Counting Chamber Slides.

### Single-cell library construction and sequencing

Approximately 20,000 cells were captured in droplet emulsions and the mRNA of single-cell libraries were constructed according to the DNBelab C Series Single-Cell Library Prep Set (MGI, Shenzhen, China) (Han et al., 2022). In brief, single-cell suspension were subjected to a series of progress, including droplet encapsulation, emulsion breakage, mRNA captured bead collection, reverse transcription, and cDNA amplification and purification, to generate barcoded libraries. Indexed sequencing libraries were established based on the instruction’s protocol. The quality supervision of libraries was implemented with a Qubit ssDNA Assay Kit (Thermo Fisher Scientific, Waltham, USA). Libraries were further sequenced by the DNBSEQ sequencing platform at the China National GeneBank.

### Single-cell RNA sequencing data processing

The raw single-cell sequencing data were processed by DNBelab C Series scRNA analysis software. Reads were aligned to the reference genome (Ensemble assembly: Sus scrofa11.1) to generate a digital gene expression matrix by STAR (Wang et al., 2022). The quality control parameters involving in gene counts per cell, UMI count per cell and % mitochondrial genes were stipulated. Cells genes were expressed in less than three cells, and cells were removed on the basis of detected genes number with a minimal of 200. Mitochondrial gene expression was set at a threshold of 5% for per cell. For each library, the doublet was removed using DoubleFinder with the default parameter (Wang et al., 2022). Then, the aligned reads were filtered to obtain cell barcodes and unique molecular identifiers (UMI) for gene-cell matrices, which were used for downstream analysis.

### Identification of cell clustering and cell type

After the initial DNBelab C Series scRNA analysis software processing, the cells were pre-processed and filtered. The data were normalized per sample using NormalizaData with default options and highly variable genes were calculated by FindVariableFeatures and then elected based on their average expression and dispersion. The cell cluster was presented with the standard integration process of P value < 0.01 through the “FindClusters” function described in Seurat (Wang et al., 2022). The cell-types in each cell cluster were identified with enriched expression using “FindAllMarkers” function in SCSA with default parameters, together with canonical cell-type markers from extensive reported literature on pig and human skin. Gene with |log2FC| > 0.25 and adjusted p-value < 0.05 were considered marker genes. And subsequently the cell cluster was visualized with t-SNE plot.

### Identification of DEGs among multiple compared groups and GO/KEGG enrichment analysis

We used the FindMarkers function in Seurat to confirm skin related DEGs between CH-back and CH-head, CH-back and CH-ear, CH-back and CH-shoulder, CH-back and CH-abdomen, CH-back and CH-leg, CH-shoulder and CH-head, CH-shoulder and ear, CH-shoulder and CH-abdomen, CH-shoulder and CH-leg, CH-head and CH-ear, CH-head and CH-abdomen, CH-head and CH-leg, CH-leg and CH-abdomen, CH-leg and CH-ear, and CH-abdomen and CH-ear for each cluster. DEGs of 15 compared groups were identified with |log2FC| > 0.25 and adjusted p-value < 0.05. In the global clusters, Gene ontology (GO) analysis was implemented with the Dr. Tom platform of BGI. Kyoto Encyclopedia of Genes and Genomes (KEGG) analysis, which was also performed with the Dr. Tom platform of BGI, further identified gene biological function including signal transduction pathways, metabolic pathways and so on in dermal cell populations.

### Cross-species comparison for skin cell atlas in pigs, humans and mice

Published skin single-cell datasets of humans (Solé-Boldo et al., 2020; Zou et al., 2021) and mice (Joost et al., 2020; Ko et al., 2022) were download from GEO with a 10 X sequencing platform. The count matrices of the three species were integrated for clustering using the Seurat R package with standard process for interspecies skin cell atlas analysis. The expressed genes that were orthologous were kept in the three species. The comparison of cell numbers and UMI count matrices was obtained for pigs, humans and mice. And the cell types were annotated by cell-type marker genes identified in this study.

### Pseudotime analysis

The cell pseudotime trajectory was constructed using R package Monocle2 (Trapnell et al., 2014). This method arranges these cells on a trajectory that describes the complete differentiation process as a quasi-time sequence of these cells through the asynchronous nature of each cell in the differentiation process.

### Cell-cell communication inference

To understanding global communication among the cell types of pig skin, we used the R package CellChat (v1.0.5) (Trapnell et al., 2014) with ligand-receptor interactions for visual intercellular communications from scRNA-seq data. As the database covers the human species, we select these pig genes according to their homologous with humans. CellChat implement some visualization methods, including the interaction number, interaction weight, communication patterns of incoming river plot, communication patterns of outgoing river plot, functional pathways, structural pathways, chord plot, circle plot, hierarchy plot and ligand-receptor of contributions.

### Targeted transcription factors interaction among cells

The transcription factor (TF) list for pig species was downloaded from the AnimalTFDB (v4.0). We identified all the TFs using motif enrichment data in cisTarget database (https://resources.aertslab.org/cistarget/), of which the “grn” module constructed a co-expression network, the “cxt” module inferred regulomes, and the “aucell” module calculated the AUC value in SCENIS (v0.11.2) (Kalucka et al., 2020). From the above data, we selected the DEGs of SMCs, ECs and FBs corresponding to TF and visualized these networks using Cytoscape software.

### Skin section

The total skin thickness from 3 pigs per breed with different sites was measured three times with a Vernier caliper at the same position and recorded. The skin tissues were fixed in a solution of 10% neutral buffered formalin and processed using routine histological procedures. Then, the sections were cut at a thickness of 5 µm using a microtome. The dermal thickness was determined using CaseViewer software according to a previous method after hematoxylin-eosin staining (Zou et al., 2022). The mean values and standard deviations were calculated.

### Immunofluorescence staining

A 5 µm-thick back skin section was incubated with primary polyclonal rabbit antibody (ABclonal, Wuhan, China) against MYH11 (1:500 dilution) and ACTA2 (1:500 dilution) overnight at 4 °C for SMCs, APOA1 (1:500 dilution) and PECAM1 (1;200 dilution) for ECs, and LUM (1;200 dilution) and POSTN (1;200 dilution) for FBs. FITC-goat anti-rabbit IgG and Cy3-conjugated goat anti-rabbit IgG were used as secondary antibodies (1:200 dilution) at room temperature for 1 hour. Then, the cell nuclei were stained with DAPI dye for 30 min. These procedures were implemented under dark conditions. Finally, these images were captured by confocal microscopy.

### Statistical analysis

Statistical testing was applied by GraphPad Prism. The data are shown as the mean ± SD for one group.

## Acknowledgements

This work supported by Chengdu Livestock and Poultry Genetic Resources Protection Center (2022) and Sichuan Science and Technology Program (2021ZDZX0008).

## Additional information

### Funding

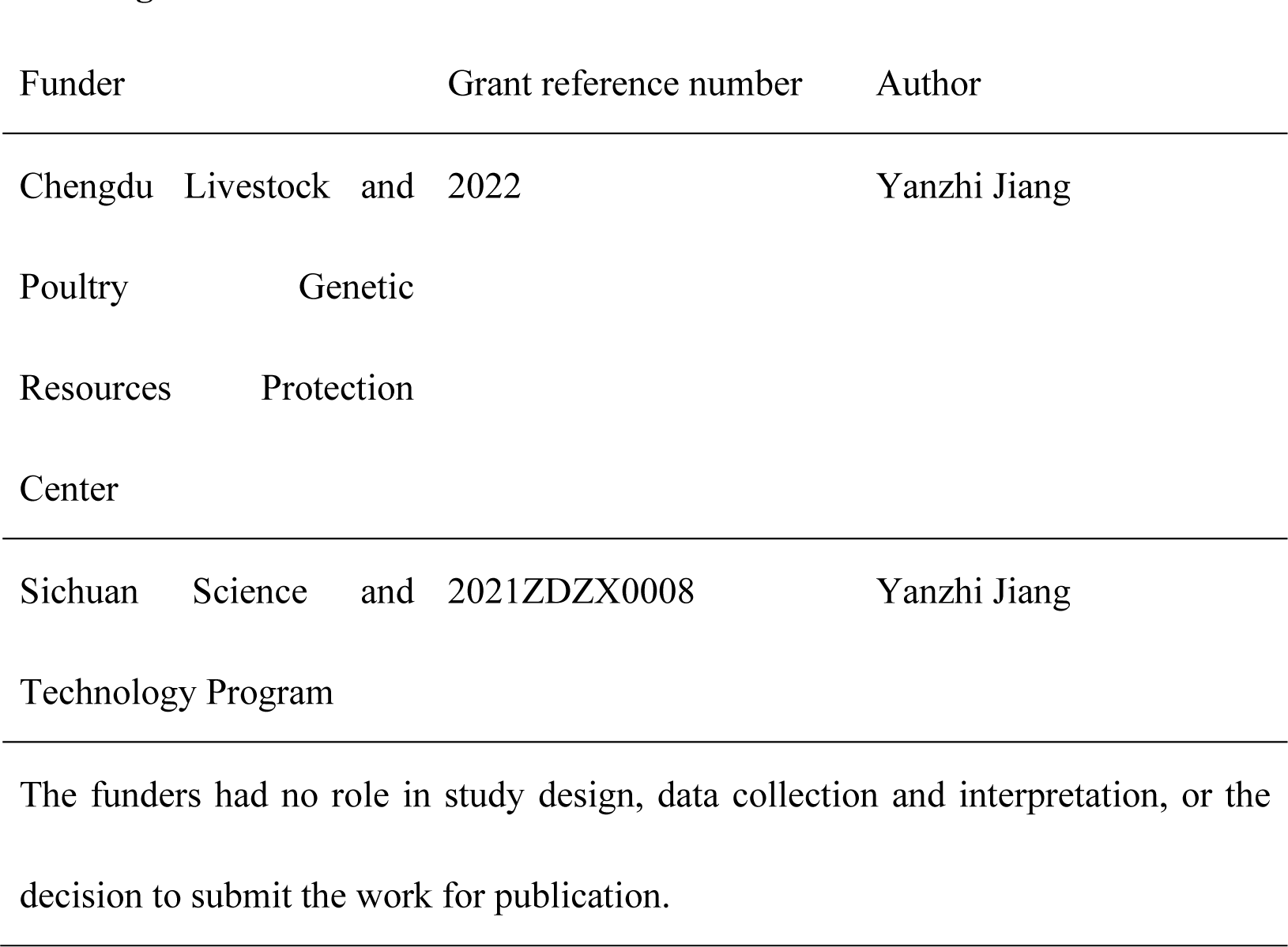

### Author contributions

Conceptualization: Q.Z., and Y.J.; Methodology: Q.Z., R.Y., Y.W., T.Z., R.S., and M.Z.; Investigation: Q.Z., R.Y., Y.L., K.F., R.F., B.P., Z.G., and X.Z.; Data analysis: Q.Z., and Y.Z.; Writing – Original Draft: Q.Z.; Writing – review and editing: Y.J., L.Z., G.T., M.L., and X.L.; Funding acquisition and Supervision: Y.J.

### Ethics

Three heads per breed aged 180 days old of both CH and LW pigs were obtained from Chengdu Livestock and Poultry Genetic Resources Protection Center. All animal experimental procedures were permitted following the Care and Use Committee of Sichuan Agricultural University (permit number: 20220219).

### Data availability

The single-cell RNA-seq data have been deposited in NCBI’s Gene Expression Omnibus database and accessible through GEO Series accession number GSE225416.

Source data files have been provided for Figures

The following dataset was generated:

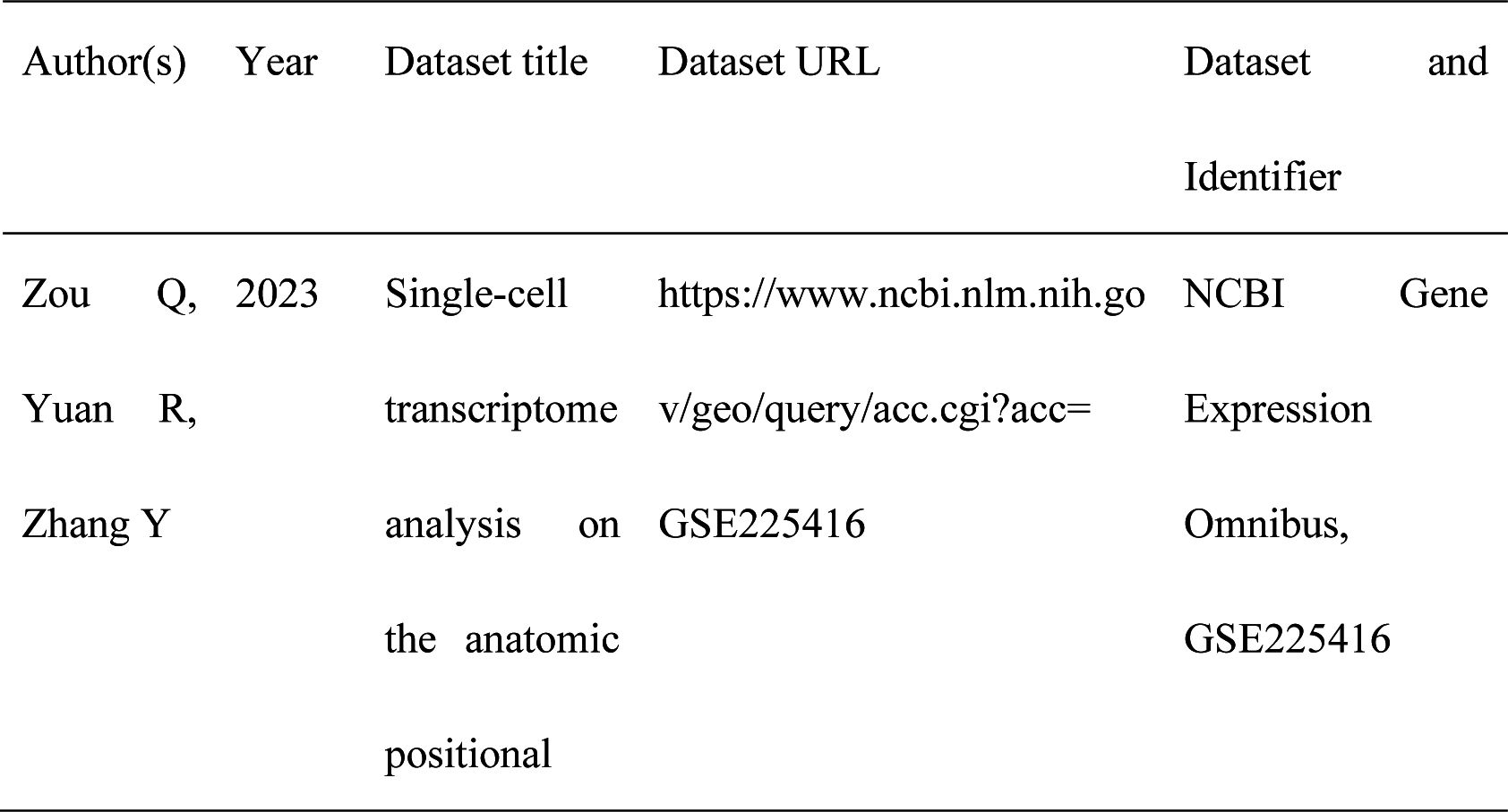
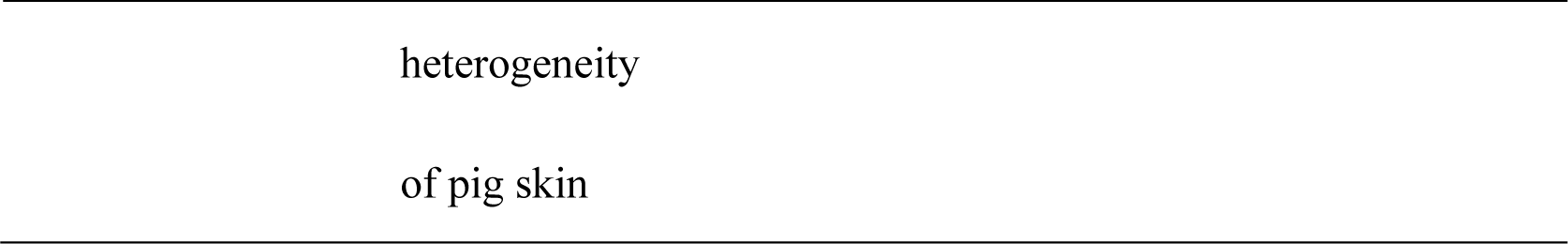

The following previously published datasets were used:

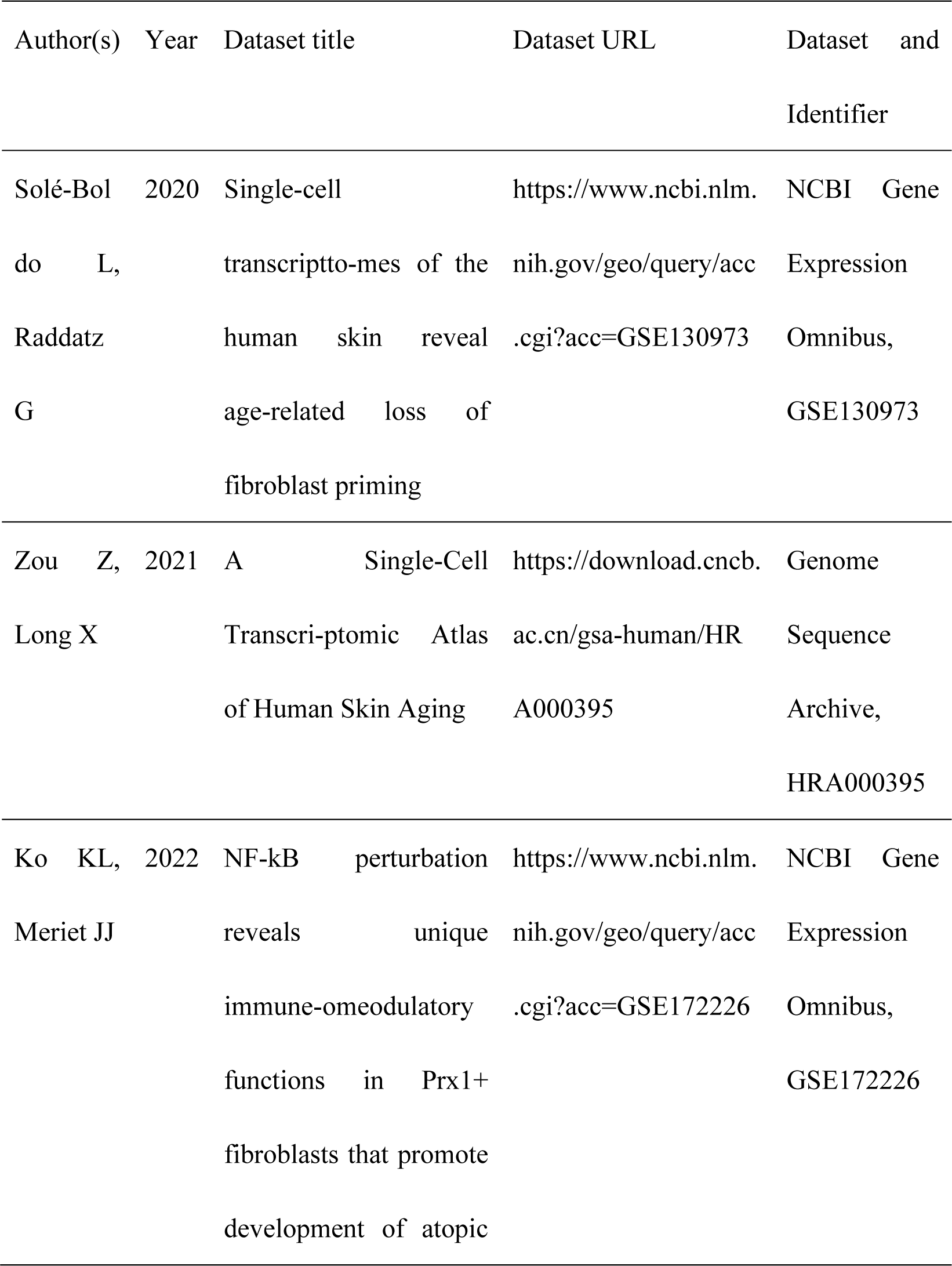
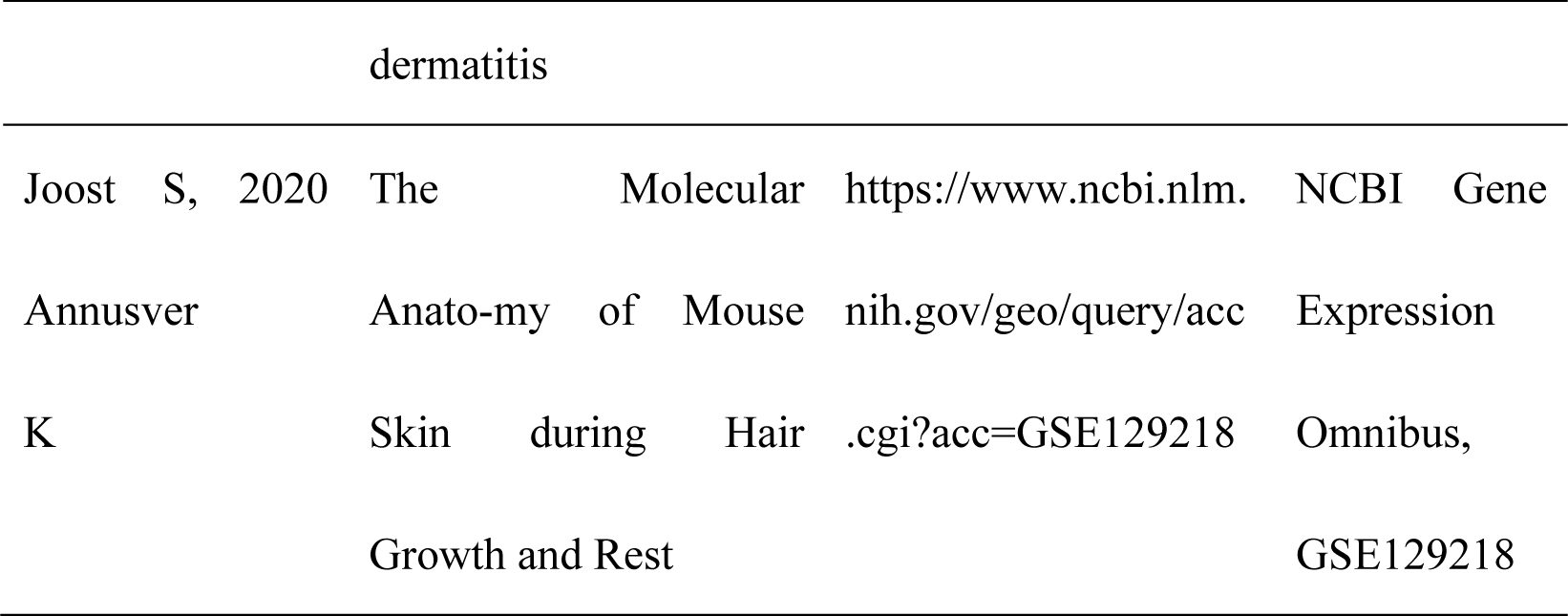

---

**Figure 1—figure supplement 1.**
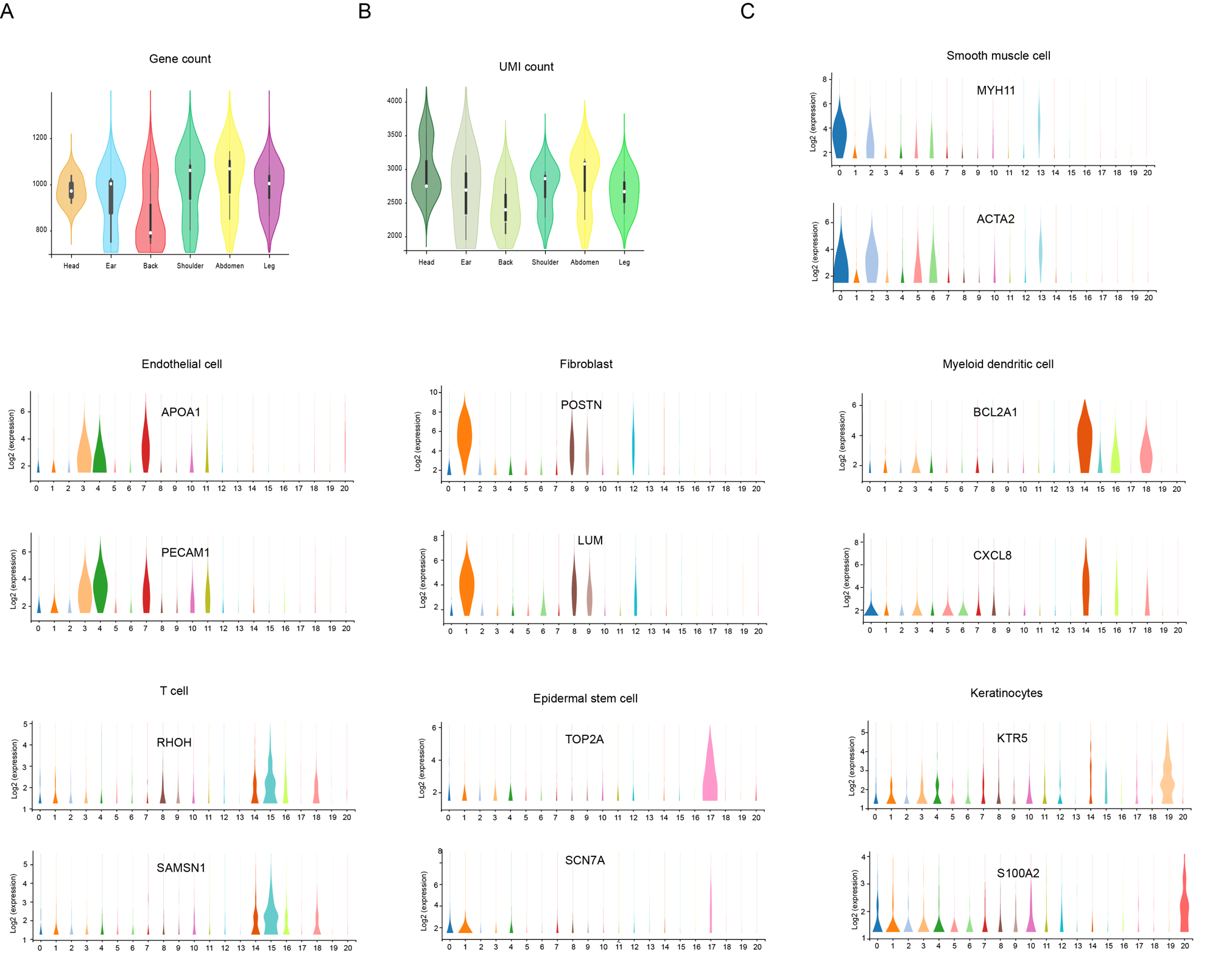
The count of genes/UMI and the expression of marker genes. (**A**) Violin plot showing the number of genes detected from different skin sites. (**B**) Violin plot showing the UMI count detected from different skin sites. **C** Violin plot showing the genes expression levels of MYH11, ACTA2, APOA1, PECAM1, POSTN, LUM, BCL2A1, CXCL8, RHOH, SAMSN1, TOP2A, SCN7A, KTR5 and S100A2 in each cell cluster of skin cells in CH pigs. **Figure 1—figure supplement 1—Source data 1.** Source data of the gene/UMI counts in figure supplement 1A and 1B

**Figure 1—figure supplement 2.**
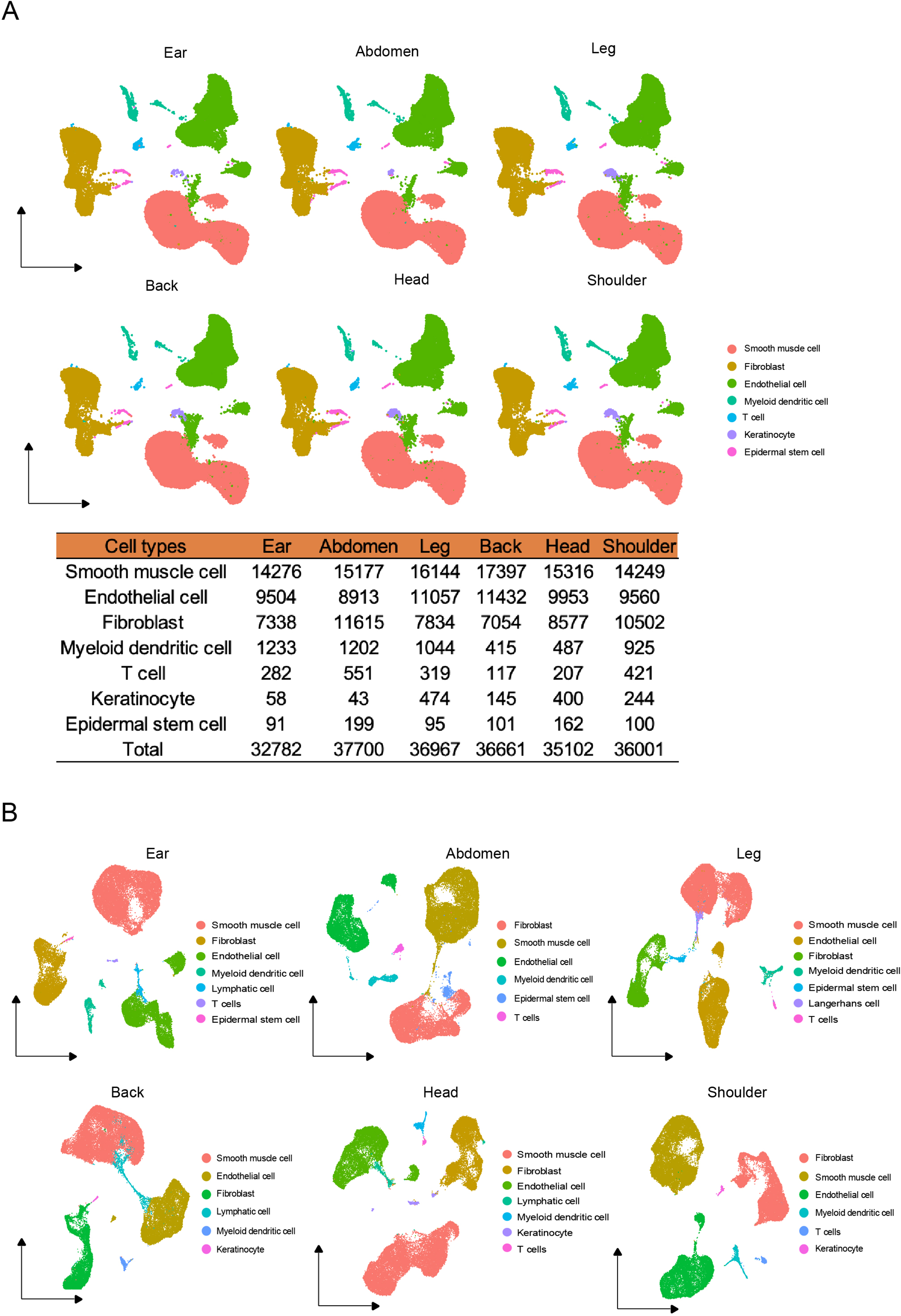
The cell types of different skin regions. **(A**) UMAP visualization of cell types from different skin regions in global CH skin cells. The number of cell types in different skin regions. (**B**) The cell types of individual skin sites by UMAP visualization.

**Figure 1—figure supplement 3.**
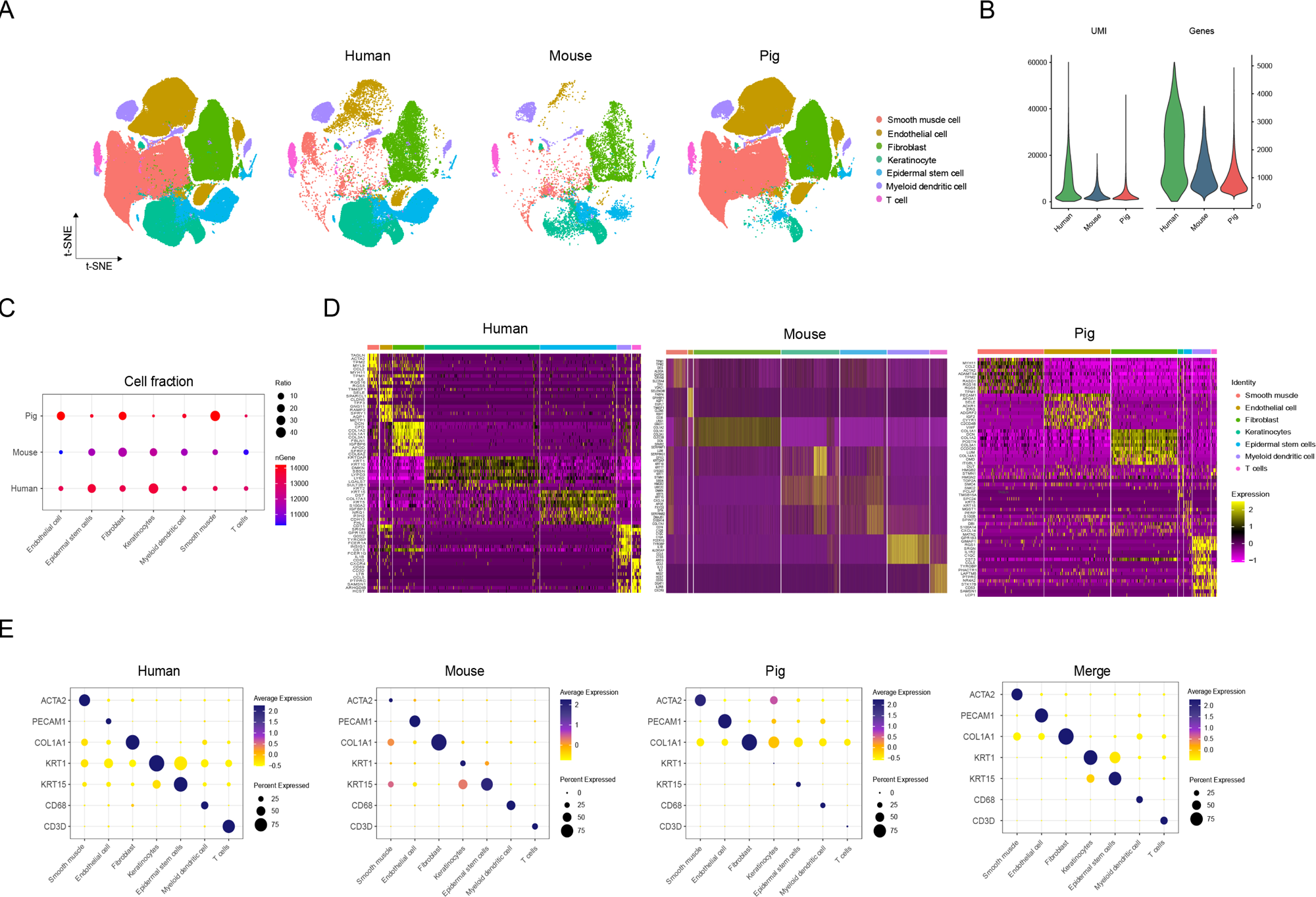
Comparison of skin cells among human, pig and mouse species. (**A**) The t-SNE plot visualization of all cell types for skin cells among humans, pigs and mice. (**B**) Violin plots showing the number of UMI and gene counts of skin cells among humans, pigs and mice. (**C**) Bubble plot representing the ratio of cell types for skin cells and the gene number among humans, pigs and mice. Color shows gene number and circle size indicates cell abundance. (**D**) Heatmap showing high expression levels of genes in each cell type of skin cells among humans, pigs and mice. Light yellow shows the genes with high expression. (**E**) Bubble plot showing the ratio and expression of marker genes in skin cells among humans, pigs and mice. Color represents genes expression and circle size indicates the percent of expressing cells.

**Figure 4—figure supplement 1.**
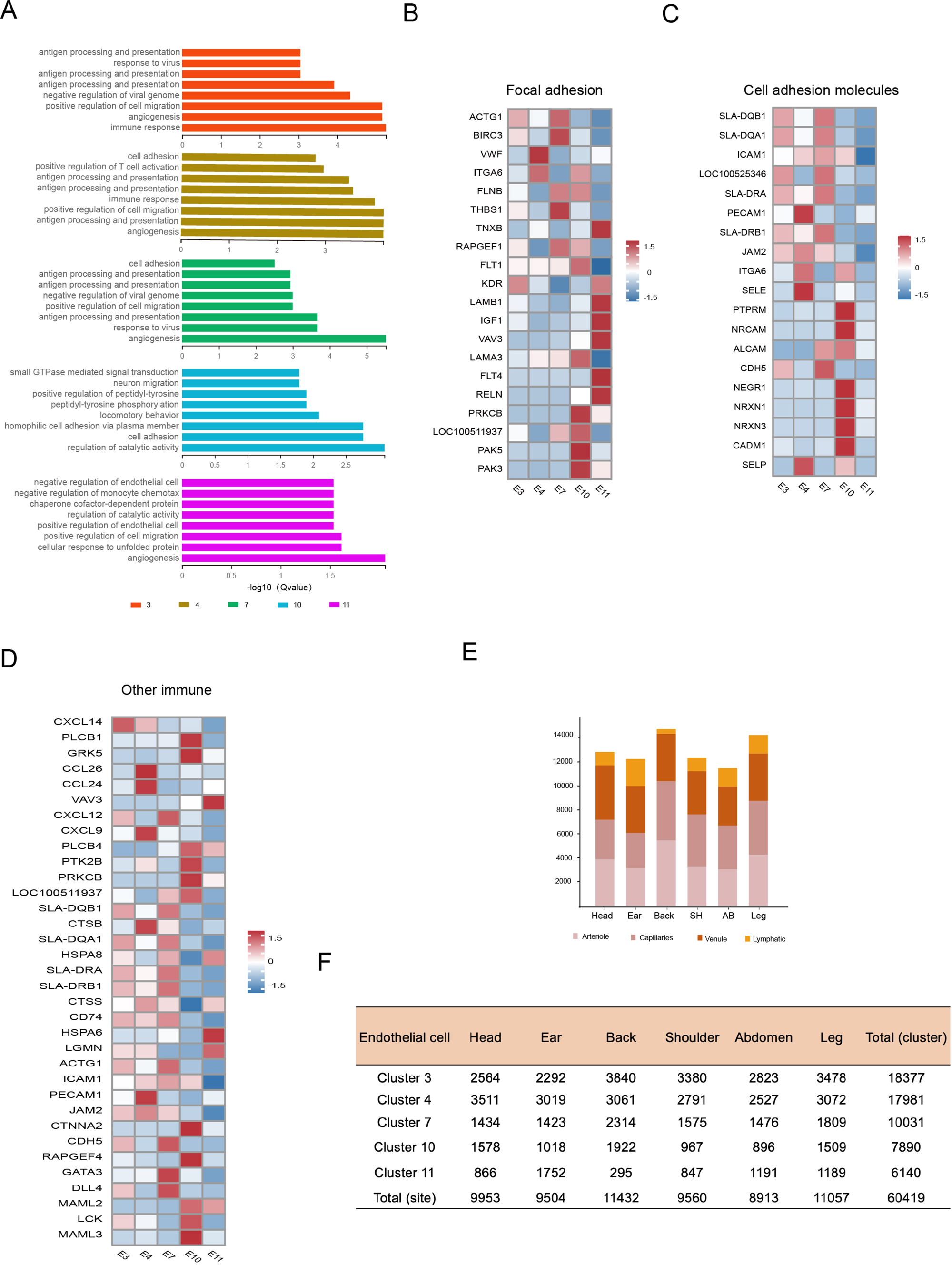
Endothelial cell heterogeneity. (**A**) The enriched GO terms pertaining to biological process for each EC subpopulations sorted by q-value. (**B**) Heatmap of gene expression for focal adhesion pathway among EC subpopulations. (**C**) Heatmap of gene expression for cell adhesion molecule pathway among EC subpopulations. (**D**) Heatmap of gene expression for other immune pathways among EC subpopulations. (**E**) The cell number of EC subpopulations in different skin regions.

**Figure 4—figure supplement 2.**
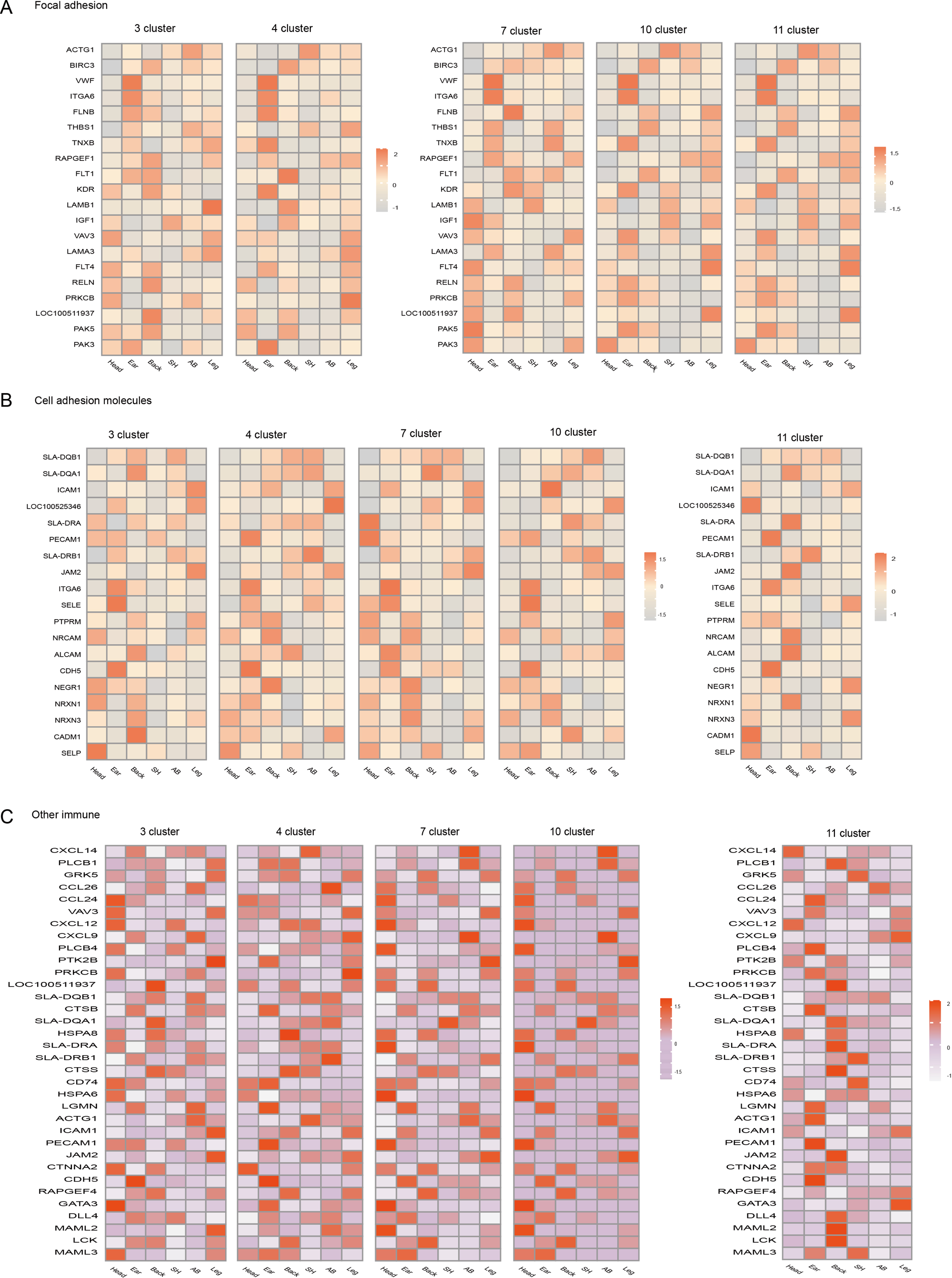
Endothelial cell heterogeneity in different skin regions. Heatmap of gene expression in focal adhesion pathway (**A**), cell adhesion molecule (**B**) pathway, and other immune pathways (**C**) among EC subpopulations of different skin regions.

**Figure 5—figure supplement 1.**
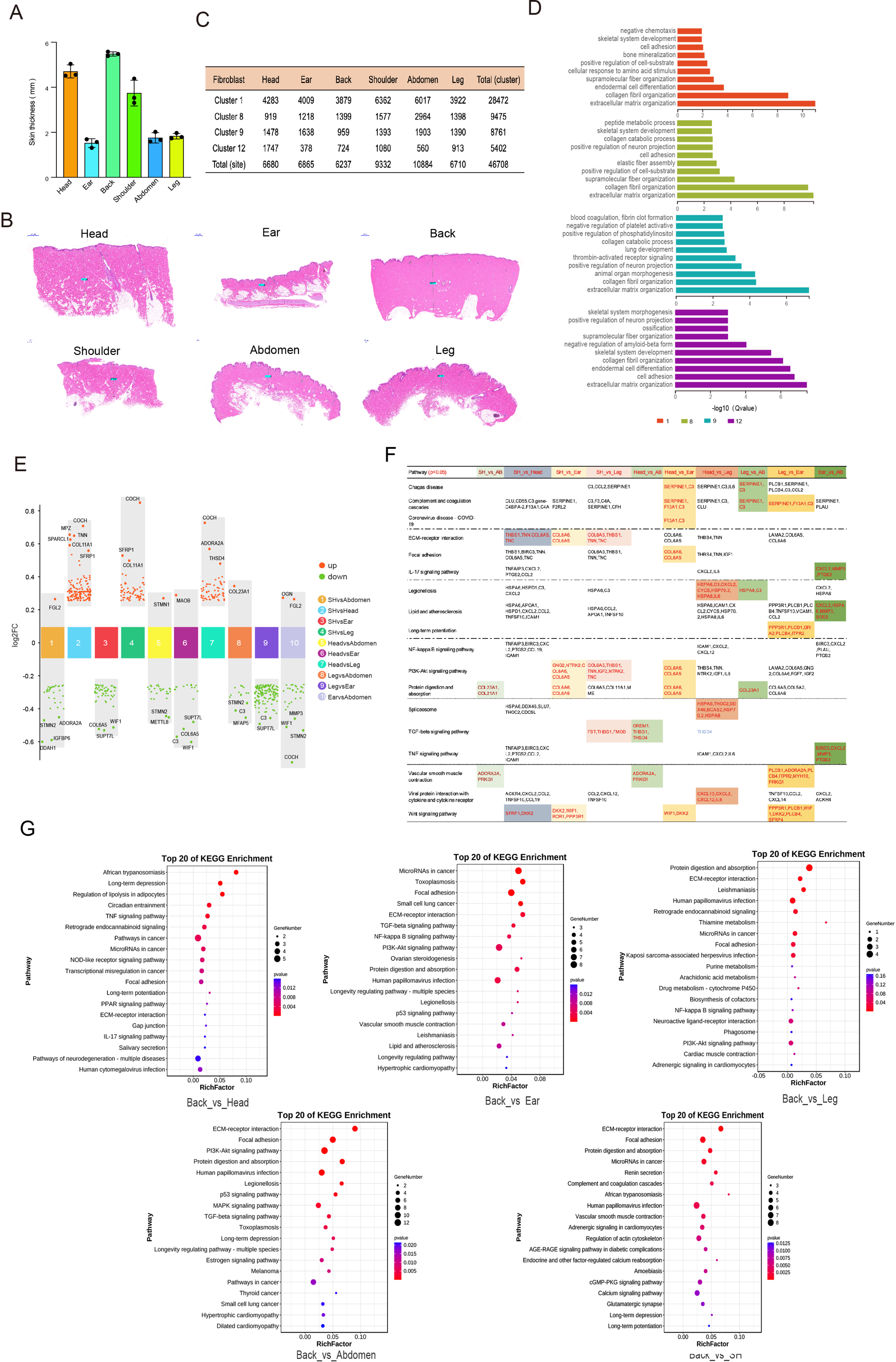
Fibroblast heterogeneity. (**A**) The skin thickness of different sites. (**B**) Skin sections of different regions. Scar bar = 1000 μm. n = 3. (**C**) The cell number of FB subpopulation. (**D**) The enriched GO terms for biological process of each FB subpopulations sorted by q-value. (**E**) Multiple volcano maps showing of DEGs of multiple compared groups. Representative genes are indicated. (**F**) Significant pathways of multi-compared groups. (**G**) KEGG analysis of back skin compared with other regions.

**Figure 6—figure supplement 1.**
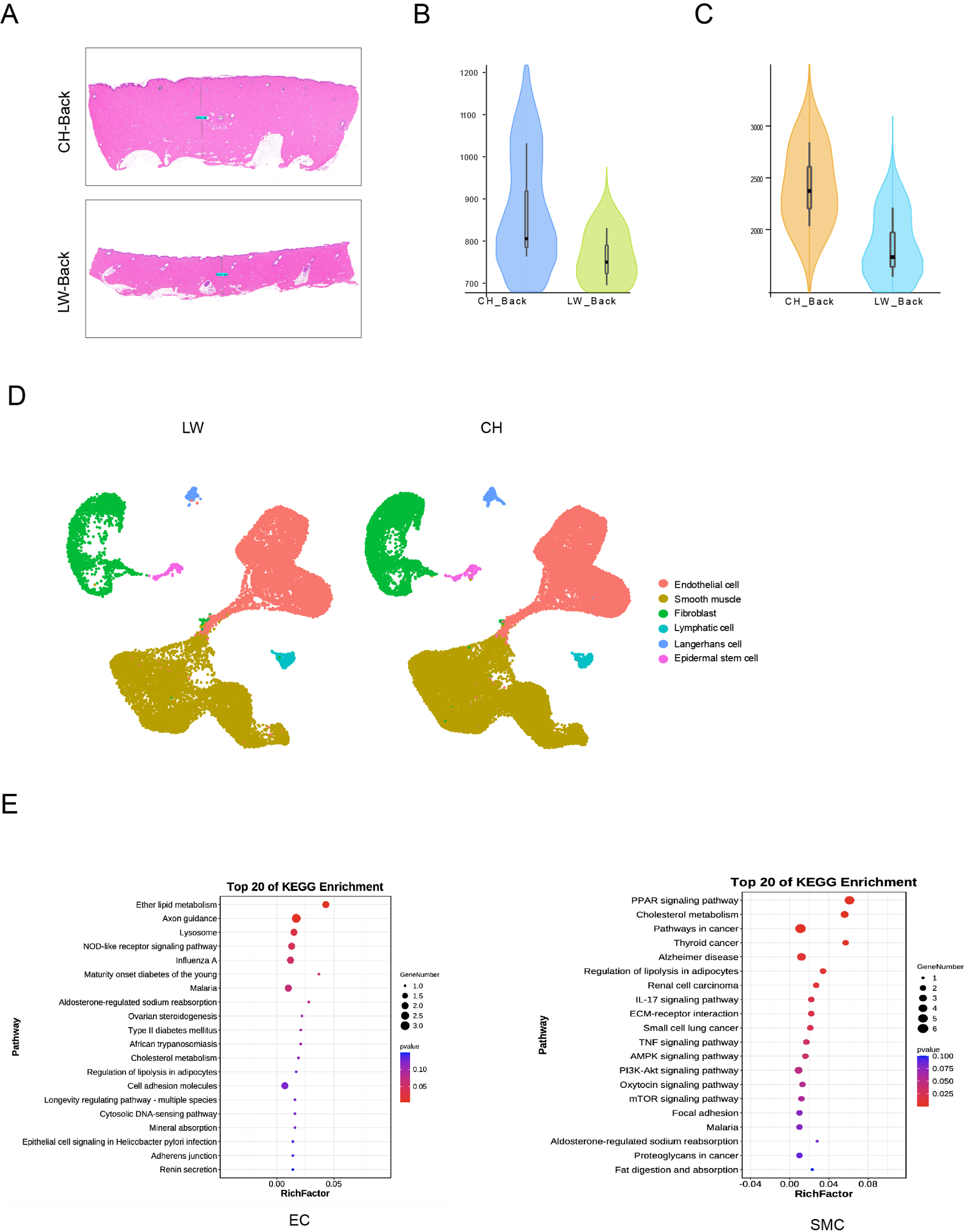
Comparison of skin cells from CH and LW pigs. (**A**) Skin sections of CH and LW pig. Scar bar = 1000 μm. n = 3. (**B**) Violin plot showing the number of genes detected from CH and LW pig skin. (**C**) Violin plot showing the UMI count detected from CH and LW pig skin. (**D**) UMAP visualization of cell types in CH and LW pig skin. (**E**) KEGG analysis of DEGs from ECs and SMCs between CH and LW pigs.

**Figure 7—figure supplement 1.**
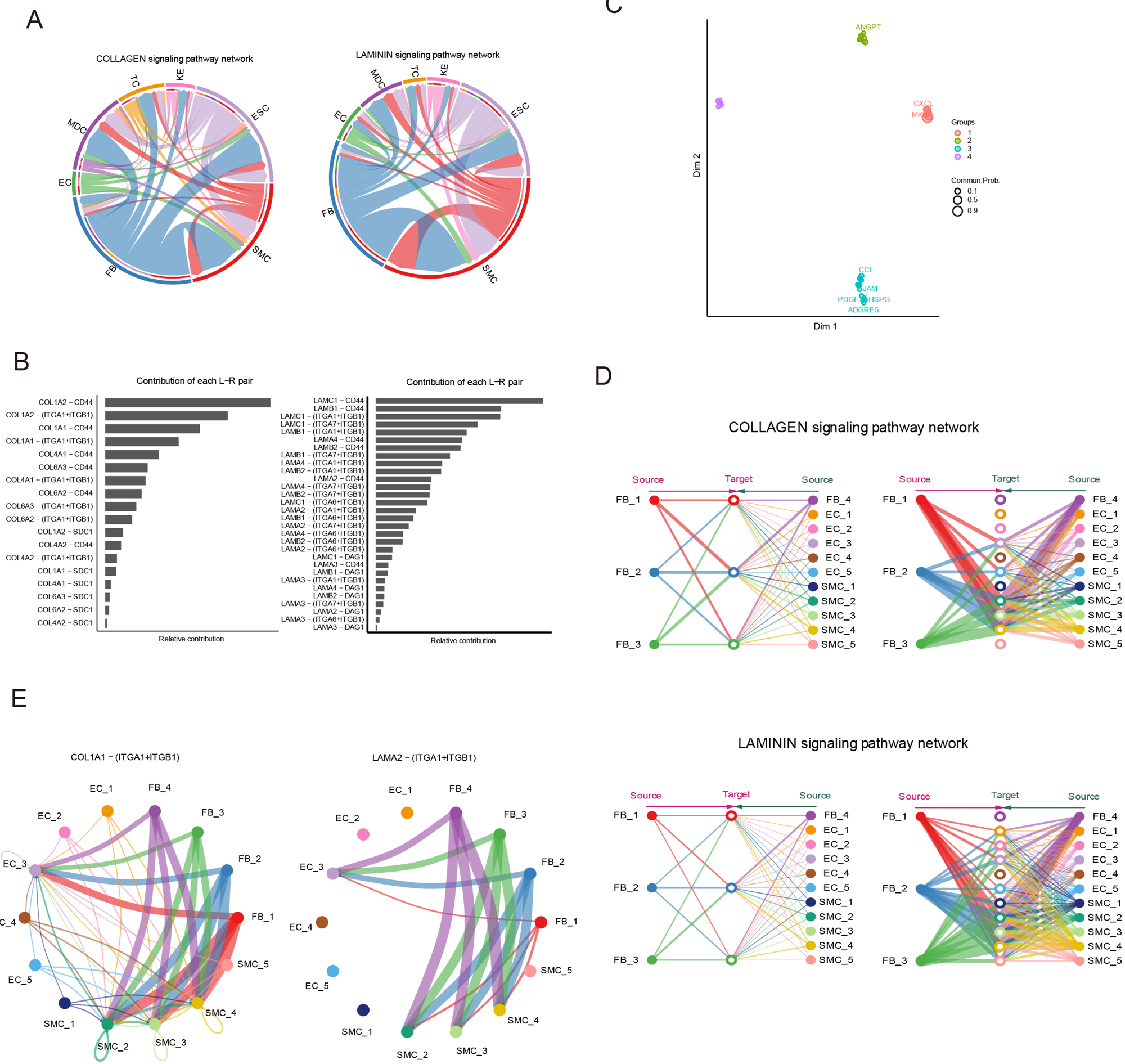
Cell communication of skin cells. (**A**) Circle plot representing the cell communication among cell types in the COLLAGEN/LAMININ signaling pathways. Edge width represents communication probability. (**B**) Relative contribution of each ligand-receptor pair of the COLLAGEN/LAMININ signaling pathway. (**C**) Projecting signaling pathway in a two-dimensional manifold based on their structural similarity. (**D**) Hierarchical plot showing the intercellular communication network of subpopulations of SMCs, ECs and FBs for COLLAGEN/LAMININ signaling pathways. Circle sizes represent the number of cells and edge width represents communication probability. (**E**) Circle plot representing the cell communication of the central ligand-receptor pair of the COLLAGEN/LAMININ signaling pathway in subpopulation of SMCs, ECs and FBs.

